# Modeling gene regulation in response to wounding: temporal variations, hormonal variations, and specialized metabolism pathways induced by wounding

**DOI:** 10.1101/2020.07.15.204313

**Authors:** Bethany M. Moore, Yun Sun Lee, Erich Grotewold, Shin-Han Shiu

**Author notes:** Department of Botany, University of Wisconsin-Madison, WI.

## Abstract

Plants respond to wounding stress by changing gene expression patterns and inducing jasmonic acid (JA), as well as other plant hormones. This includes activating some specialized metabolism pathways, including the glucosinolate pathways, in the case of *Arabidopsis thaliana*. We model how these responses are regulated by using machine learning to incorporate putative cis-regulatory elements (pCREs), known transcription factor binding sites from literature, *in-vitro* DNA affinity purification sequencing (DAP-seq) and DNase I hypersensitive sites to predict gene expression for genes clustered by their wound response using machine learning. We found temporal patterns where regulatory sites and regions of open chromatin differed between clusters of genes up-regulated at early and late wounding time points as well as clusters where JA response was induced relative to clusters where JA response was not induced. Overall, we identified pCREs that improved model predictions of expression clusters over known binding sites. We discovered 4,255 pCREs related to wound response at different time points and 2,569 pCREs related to differences between JA-induced and non-JA induced wound response. In addition, pCREs found to be important at different wounding time points were mapped to the promoters of genes in a glucosinolate biosynthesis pathway indicating regulation of this pathway under wounding stress. Finally, we experimentally validated a predicted cis-regulatory element, CCGCGT, showing that knock-out via CRISPR-Cas9 reduces gene expression in response to wounding.

## Introduction

Plants cope with many environmental stresses by reprogramming their pattern of gene expression to trigger chemical and physiological responses (Bostock et al., 2014). These stress responses are essential to plant survival in their respective niches and are optimized for a plant’s particular environment (Bostock et al., 2014). Gene expression reprogramming is a complex process that involves multiple levels of regulation. At the DNA sequence level, short stretches of DNA (regulatory elements) are recognized and bound by transcription factors that can activate or repress gene expression (Zou et al., 2011). Beyond the level of DNA sequence, chromatin structure can impact whether a regulatory element is accessible to a transcription factor.

Chromatin structure can be modified based on signals stress response signals (Asensi-Fabado et al., 2017). Finally, reprogramming can also occur by modifying (Glisovic et al., 2008) or turning over (Hutvagner and Simard, 2008) messenger RNA.

Stress responses change over time, adding an additional level of temporal complexity to transcriptional response. For example, after an initial response, genes that are turned on may act to turn on or off other genes, resulting in cascading effects. This type of gene expression reprogramming mechanism is beneficial when different responses are needed at different times. For example, response to wounding stress in plants changes over time as the plant first needs to recognize damaging agents, then respond by sending various hormone signals, and ultimately repair the wound (Ikeuchi et al., 2017). This means that stress responsive genes may be regulated differently depending on when they are expressed.

The production of various hormone signals allows plants to coordinate their response to different stresses because the interactions of certain hormones can regulate a specific response from the plant by changing the expression of certain genes. For example, response to wounding stress involves several hormones, with the most ubiquitous signal being jasmonic acid (JA) (Howe and Jander, 2008). After wounding, JA levels increase and bind to JAZ repressor proteins, which allows MYC2 transcription factors to become active (Chung et al., 2008). MYC2 transcription factors then activate wounding responses, such as JA biosynthesis, to amplify the JA signal and activate other defensive processes (Chung et al., 2008). Additional hormones interact with JA to moderate wounding response. For example, while JA induces the expression of certain wounding response genes, ethylene simultaneously represses the expression of these genes at the damaged site in order to make sure the correct spatial response pattern is produced (Rojo et al., 1999). Ethylene also works in a synergistic fashion with JA to fine-tune wounding response by inducing the expression of proteinase inhibitor genes (O’Donnell et al., 1996) and by activating ERF1, another transcription factor that triggers defense responses (Lorenzo et al., 2003). Abscisic acid (ABA), which is induced in response to many abiotic stresses, is also induced by wounding (León et al., 2001). While ethylene, ABA, and JA rapidly respond to wounding, other hormones such as auxin and cytokinin, start to accumulate around 12 hours after wounding and are involved in signaling for the expression of genes that work to repair the wound (Ikeuchi et al., 2017). While a great deal is known about hormone signaling in response to wounding, it is unclear what other regulatory mechanisms are involved in response to wounding and how these mechanisms interact with hormone signals. In particular, regulatory mechanisms for wounding responses not directly regulated by JA are less well understood.

Wounding can also induce the production of specialized metabolites that can deter further stress. For example, after wounding stress, *Arabidopsis thaliana* activates glucosinolate pathways. These glucosinolates and the bioproducts generated from their degradation affect the plant’s interactions with biotic stresses, such as microbes and herbivores (Yan and Chen, 2007). Additionally, mutants with decreased glucosinolate levels show greater susceptibility to the necrotrophic fungus *Fusarium oxysporum* (Tierens et al., 2001). Glucosinolate production is regulated by JA, salicylic acid (SA), and ethylene (ACC). These hormones work together to modulate glucosinolate levels in response to stress, by activating MYB and DOF transcription factors (Yan and Chen, 2007). Additionally, glucosinolates can be divided into different types, such as indole or aliphatic glucosinolates, and these types may be induced by various stresses and regulated in different ways (Yan and Chen, 2007). While specific transcription factors have been shown to turn on glucosinolate biosynthesis (Frerigmann and Gigolashvili, 2014), the regulatory elements or chromatin structure of how and when these transcription factors bind has not been resolved.

Here, we assessed the extent of divergence in gene expression among various time points following wounding by correlating wounding data with other types of stress or hormone treatment. By using a time course data set, where transcriptional response were recorded over a 24-hour period (Kilian et al., 2007), we captured differences in differential gene expression and the regulatory elements required to regulate this transcriptional response. Because most regulatory elements occur 1000 bp upstream in the promoter region of the gene (Weirauch et al., 2014; Yu et al., 2016), we focused on this region to identify putative *cis*-regulatory elements (pCREs). In addition, by clustering wound-responsive genes into groups based on whether or not they also respond to JA, we were able to single-out differences between JA and non-JA regulatory mechanisms in regard to wounding. By using a timecourse study, we identified important regulatory elements glucosinolate biosynthesis from tryptophan, which is induced by wounding. Finally, by using machine learning modeling, we were able to identify the most important regulatory sequences for each time point and experimentally show that a previously unknown element affects wounding response. The goals of this study were to uncover the cis-regulatory code involved in regulating temporal responses to wounding stress, to see how wounding stress independent of the wound-induced hormone JA is regulated, and finally to understand how certain specialized metabolism pathways are regulated.

## Results

### Transcriptional response to wounding varies functionally across time points

To understand how transcriptional response to wounding varies across time points, we used expression data downloaded from TAIR, in which a range of abiotic stress treatments (seven in total, including wounding) were applied to 18 day old *A. thaliana* seedlings (Kilian et al., 2007). Samples were harvested at multiple time points after treatments ranging from 15 minutes to 24 hours after treatment. Control samples were performed in parallel to exclude circadian effects (see **Methods**, Kilian et al., 2007). We identified genes that were up- or down-regulated at time points ranging from 15 minutes to 24 hours after wounding (diagonal values; **Figure 1A**) and how frequently the same genes were differentially expressed in these different time points (lower triangle; **Figure 1A)**. We found a cascading effect, where the majority of genes up-regulated at 15 and 30 minutes after wounding are still up-regulated at one hour (63% and 70% respectively), but by three hours <25% of those genes were still up-regulated (**Figure 1A, Table S1**). Consequently, the genes up- or down-regulated at later time points tended to be different from those differentially expressed earlier, with the genes responsive at 12 and 24 hours after wounding having the least amount of overlap with genes from previous wounding time points (**Figure 1A, Table S1**). Thus, different time points after wounding have overlapping but distinct sets of genes which are up- or down-regulated, suggesting temporal variation in how wound response is regulated.

**Figure 1.**
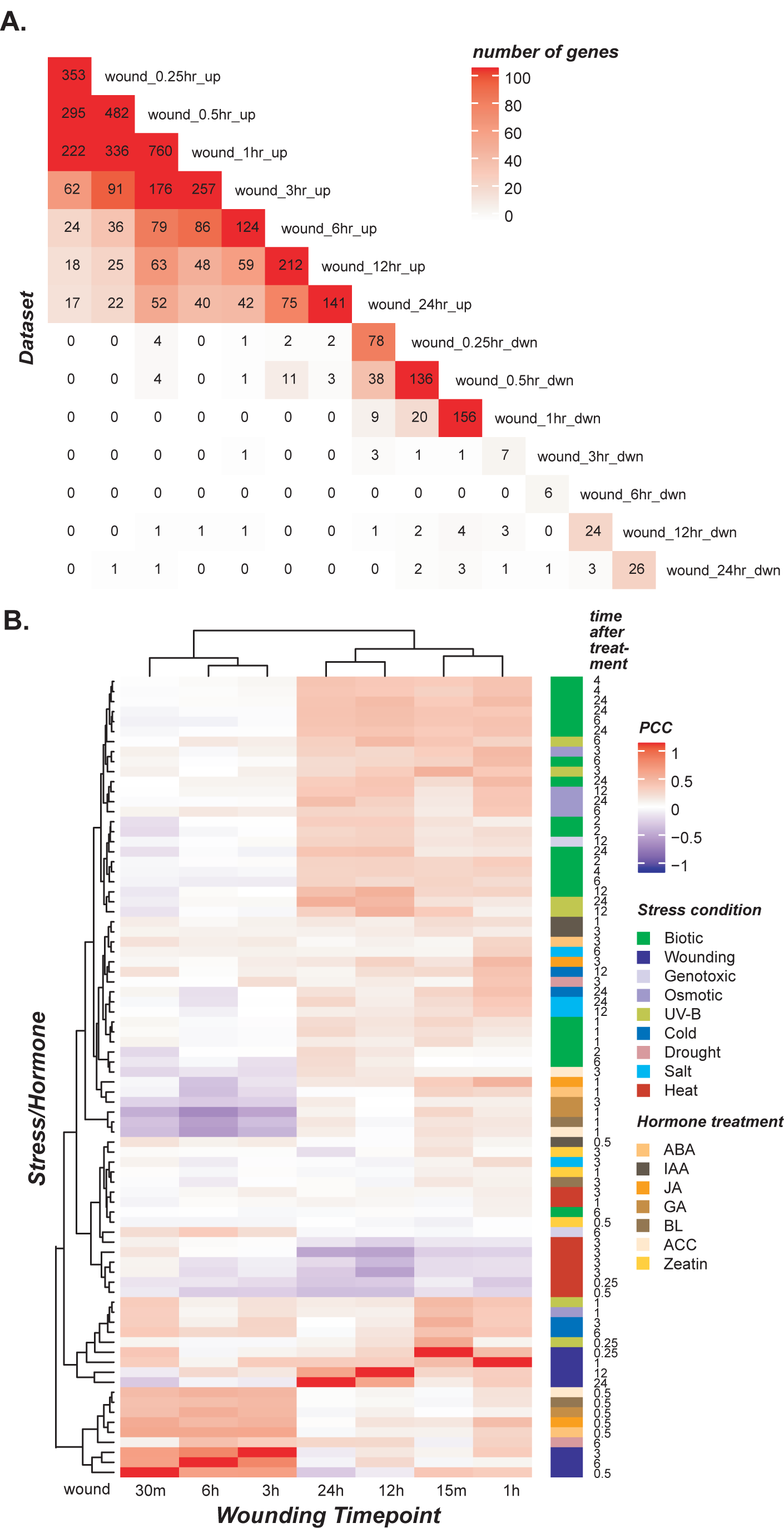
Gene expression correlation across stress and hormone data sets and the overlap of wound and JA differentially expressed genes. A. Heatmap showing the number of genes overlapping in each wounding time point cluster. The order of rows and columns are the same, based on time point. Number of genes range from 0 (white) to 760 (red) and actual value is printed in the heatmap. B. Heatmap of Pearson’s correlation coefficient (PCC) between data sets based on the log2 fold change between treatment and control. PCC values in heatmap range from 1 (red) to -1 (blue). The order of the rows and columns are the same, which is based on hierarchal clustering. The stress and hormone treatments are labeled by color and stress time point is labeled on y-axis.

To determine how response to wounding differs from response to other environmental conditions, we measured how similar the pattern of differential gene expression was between different wounding time points and other abiotic stress, biotic stress, and hormone treatments (also downloaded from TAIR, see **Methods**). The Pearson’s correlation coefficient (PCC) was used to compare the log_2_ fold change values across genes between wounding and other stress/hormone treatments and then hierarchal clustering was used to find conditions where differential gene expression was most similar (**Figure 1B**). We found that gene expression patterns 30 minutes, 3 hours, and 6 hours after wounding clustered together, 24 hours and 12 hours after wounding cluster together, and then 15 minutes and 1 hour after wounding cluster together. Because wounding can be due to abiotic (i.e. wind damage) or biotic (i.e. insect chewing) stresses, to better understand this pattern, we looked at how wounding time points correlated with stress response.

Are patterns of wound stress response unique to abiotic or biotic stresses, or do they represent a general stress response? Overall, we found wound response correlated with both abiotic and biotic stress patterns (**Figure 1B**). Early patterns of wounding DGE (15, 30 minutes or 1 hour after wounding) correlated more strongly with those of early abiotic stress response compared to later wounding time points (12 or 24 hours after wounding). For example, DGE responses 15 minutes after wounding and 15 minutes after UV-B light treatment were most highly correlated with each other with a PCC of 0.51 (for PCC results, see **Table S2)**. Gene expression patterns at 30 minutes and 1 hour after wounding were also more similar to early responses (30 minutes to 3 hours) under certain abiotic stresses, such as cold, UV-B, osmotic, and genotoxic stress (**Figure 1B, Table S2**). Additionally, early DGE 15 minutes and 1 hour after wounding were more similar to each other (PCC= 0.39) and to 30 minutes after wounding (PCC= 0.33 and 0.30 respectively) than to later wounding time points (for PCC results, see **Table S2**). The DGE response from the 12 and 24 hour time points also had a high correlation to the DGE response at 12 or 24 hours after UV-B or osmotic treatment, showing late wound response is more similar to other late abiotic stress responses than to early wound response.

Thus, transcriptomic responses were more similar among comparable time points between treatments than largely differing time points within a particular stress. This indicates that temporal patterns can impact gene expression more than the type of abiotic stress, and that wounding can elicit a similar response to other types of abiotic stress.

The biotic stresses included pathogens *Pseudomonas syringae* and *Phytophthora infestans*, as well as pathogen-derived elicitors Flagellin (bacterial), necrosis-inducing *Phytophthora* protein (oomycete), and Hairpin Z (bacterial). When observing wounding response patterns in relation to biotic stress 15 minute, 1, 12, and 24 hour time point responses all correlate with the different types of biotic stress listed above, the 12 and 24 hour time point DGE responses have a higher correlation to (PCC range from 0.35 to 0.47) the DGE responses to *P. infestans* than any other wounding time point (**Figure 1B, Table S2**). Thus, wounding response is similar to both abiotic and biotic stress, however later time points are more highly correlated with biotic stress. *P. infestans* is a necrotrophic oomycete that creates extensive tissue damage in the plant, which may be similar to wounding damage, and the later time points of both stresses may be most similar because of similarities in immune response and recovery. It is interesting that wound DGE response at 3 and 6 hours after wounding do not correlate with biotic stress response (PCC range -0.09-0.07, **Table S2**). One hypothesis is that initial response to wounding triggers some of the same pathways involved in response to other biotic stresses and the late response to wounding triggers pathways involved in recovery from other biotic stresses, but the middle time points are involved in separate functions from biotic stress. This could explain similarities in DGE response seen between early and late time points.

Hormonal responses are also triggered by wounding, including JA, ABA, and ACC (ethylene), and correlate with wound response. Also, certain hormones, including IAA, ABA, and JA have a positive relationship where the hormones up- or down-regulate the same genes (Goda et al., 2008). While DGE 15 minutes after wounding was not similar to DGE after hormone treatment, by 30 minutes after wounding, DGE was similar to DGE 30 minutes after treatment with five different hormones (ABA, ACC, brassinosteroid (BL), gibberellic acid (GA), and JA, PCCs ranging from 0.37-0.52; **Figure 1B, Table S2**), indicating a strong temporal component to how genes are expressed, and the initial response to wounding may involve multiple hormones. The DGE responses at time points at 3 and 6 hours after wounding were more similar to the DGE response to plant hormones at 30 minutes including ABA, ACC, BL, GA, and JA (PCC range from 0.54-0.39), than most other wounding time points with the exception of 30 minutes after wounding (PCC range from -0.04-0.21; **Figure 1B, Table S2**).

Similarities between several hormonal responses and response to wounding at mid-range time points indicate that many stress-responsive hormones may still be involved in wound response even after 6 hours. Finally, 12 and 24 hours after wounding, transcriptomic responses show little correlation with DGE responses to the hormones correlated with earlier time points, with the highest correlation to DGE response to JA treatment after 3 hours (PCC = 0.26 and 0.14, respectively, **Table S2**). This indicates that the later responses to wounding may not signal stress-responsive hormones or that they do not correlate with the early transcriptomic responses to many hormone treatments. Overall, the high association of DGE patterns in early and mid-range time points after wounding to early hormone treatment DGE patterns indicates an interaction between wound response and hormone response, in which wound response likely signals various stress-related hormones, but this interaction lessens over more time after wounding as at later timepoints genes are likely involved in fixing wound damage rather than signaling.

### Modeling temporal wound response using machine learning

The temporal differences in transcriptional response to wounding described above suggest that the regulation of wounding response changes over time, with regulatory control being more similar within early time points (0.25, 0.5, 1 hours), middle time points (3 and 6 hours) and late time points (12 and 24 hours) compared to between these time points. In order to compare what regulatory mechanisms were important across different time points, we first needed to model the regulatory code of transcriptional response to wounding for time point. We used a machine learning approach to generate models of the regulatory code that could classify a gene as being differentially regulated or non-differentially regulated at a specific time point.

First, we tested how well-known sequence based regulatory information was able to model wounding response. We collected 52 known *cis-*regulatory elements (CREs) associated with JA, wounding, or insect response identified previously using experimental or computational approaches (see **Methods**; **Table S3**). To incorporate the known CRE information into a model, we mapped each putative regulatory sequence to the promoters (defined as 1 kb upstream of the transcription start site, see **Methods**) of each gene in a cluster (each cluster consisting of genes up- or down-regulated after wounding at a given time point), as well as to genes in a “null” cluster, consisted of genes that are not significantly up-regulated or down-regulated under any stress or hormone treatment. Two algorithms, Random Forest (RF) and Support Vector Machine (SVM) were used to build models for each wounding response timepoint. To measure model performance, F-measure was used which jointly considers precision, or the number of genes that were differentially expressed and were predicted as differentially expressed over the number of all of genes the model predicted as differentially expressed, and recall, or the number of genes which were differentially expressed and were predicted as differentially expressed over the number of genes which were differentially expressed (predicted or not). The F-measures for models built for each wounding time point cluster ranged from 0.67 to 0.71, scores that show our models performed better than random guessing (F-measure = 0.5) but were not perfect predictors (F-measure = 1) (for RF models: **Figure 2**, for SVM models: **Figure S1A, Table S4**). Note that three wounding response timepoints were not included in this analysis: down-regulated 3 and 6 hours after wounding because too few genes had these responses to train a machine learning model (<10) and down-regulated 12 hours after wounding category because no known regulatory elements were present in the promoters of the genes in this group.

**Figure 2.**
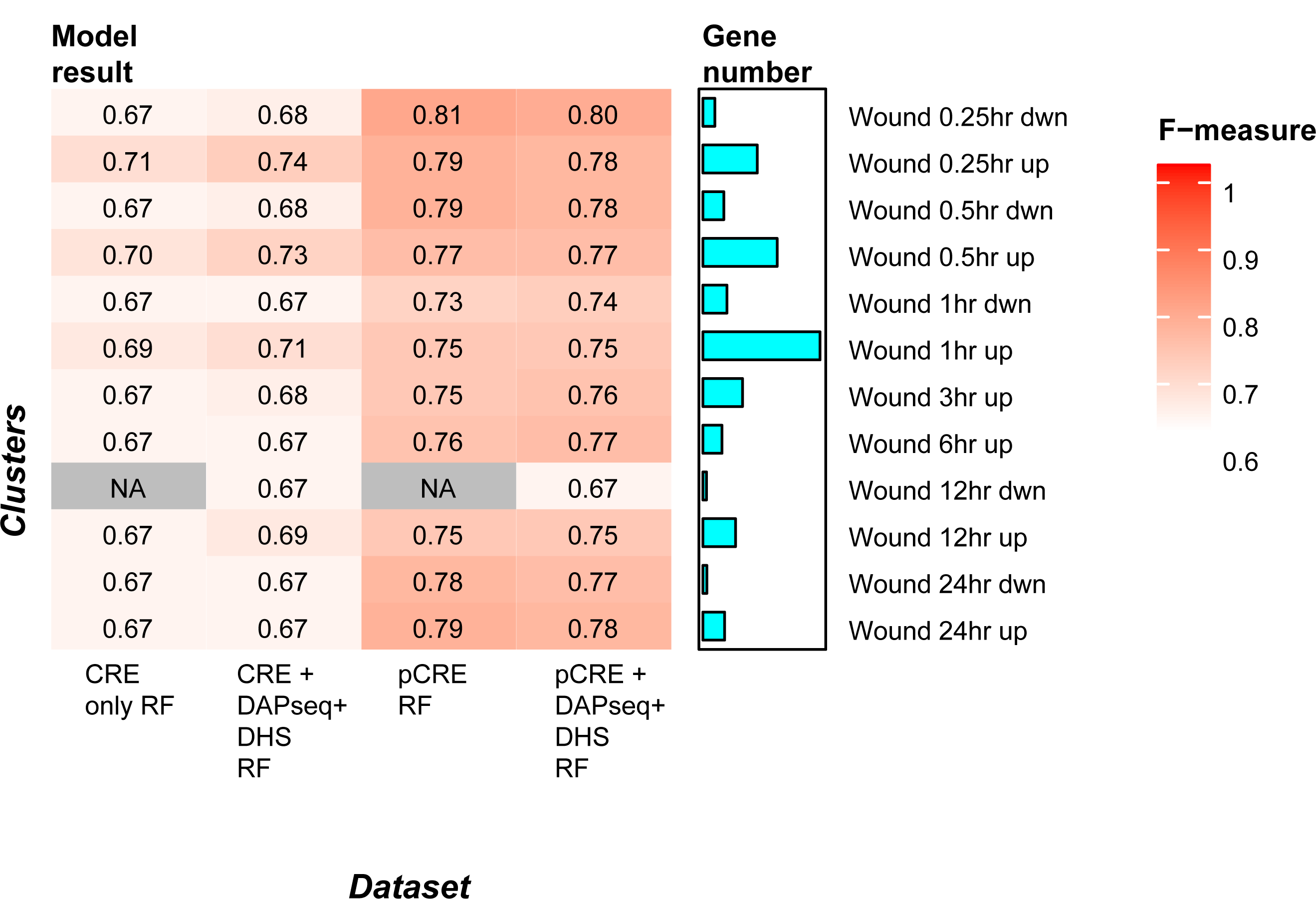
Heatmap of the F-measure for wounding time point models. Each row is a different cluster which was used to build a separate model. Each column represents the datasets used as features in the model and the algorithm used (RF= Random Forest). Known only refers to CREs found in the literature (**Table S3**). DAPseq and DHS refer to the DAP-seq and Dnase I hypersensitivity sites. FET enriched 6mer refers to the pCREs which were enriched for a specific cluster. The F-measure range is from 0.5 (white) to 1 (red), and gradient as well as actual F-measure is shown in each cell. The bar chart next to the heat map corresponds to each row/cluster and represents the number of genes in that cluster.

Second, we incorporated additional levels of regulatory information into our models. We included *in vitro* DNA binding data of 510 TFs in *A. thaliana* generated with DNA affinity purification sequencing (DAP-seq) (O’Malley et al., 2016) and information about DNase I Hypersensitive Sites (DHS) in *A. thaliana* at different developmental stages including seedling (leaf samples) and two-week old plants (flower buds) (Zhang et al., 2012). Each DAP-seq and DHS feature was considered present if its peak coordinates overlapped with the promoter region of a gene. Machine learning models trained using both known sequence and DAP-seq and DHS features performed slightly better overall than known sequence-based models alone, with the F-measure ranging from 0.66 to 0.74 (**Figure 2, Table S4**). Models for genes up-regulated in early wounding response (0.25, 0.5, and 1 hours) benefited the most from the addition of these two data sets, with a +0.03, +0.03, and +0.02 improvement in F-measure, respectively. This may be because most known CREs are known from early wound responses. Thus, more known information in the form of the DAP-seq data may improve the performance of early time point clusters more than later time points. Overall, while known sequence-based information and DAP-seq and DHS information is predictive of differential gene expression in response to wounding across time points, the models still have substantial room for improvement.

### Determining important known motifs for temporal wound response

To understand what known elements are important for driving expression at different times after wounding, we measured the importance of each feature (known CRE, DAP-seq, or DHS, see **Methods**) in each model in order to rank features in terms of how important they were for our ability to predict differential gene expression at different time points (**Table S5**). For early wound response (genes up-regulated 0.25, 0.5, and 1 hour after wounding), the most important known wound CREs identified by our models were CGCGTT (first ranked), a known regulatory element for Rapid Wound Response (RWR) (Walley et al., 2007) and CACGTG (second ranked) that is bound by some *Myc* TFs in the basic Helix-Loop-Helix (bHLH) family in response to wounding and JA treatment (Fernández-Calvo et al., 2011). Genes with the RWR elements are known to respond quickly to wounding and have a variety of functions in the downstream response, including chromatin remodeling, signal transduction, and mRNA processing (Walley et al., 2007). These functions are consistent with stress-induced transcriptional changes, where chromatin conformation is changed to modulate binding of stress-related TFs, mRNAs are modified post-transcriptionally, and signaling pathways up and down-stream to transcription are involved in response to wounding stress. Other TFs that respond to wounding stress, *Myc* 2, 3, and 4 TFs, respond to both JA and wounding, and induce other JA responsive genes, ultimately triggering defense response to herbivory (Fernández-Calvo et al., 2011). In addition to genes up-regulated 0.25 ∼ 1 hour post wounding, CACGTG, the wound response element that *Myc* TFs bind, was still important (ranked 1 or 2) among genes up-regulated 3, 6 and 12 hours after wounding, but not the RWR element. By 24 hours after wounding, the CACGTG element was no longer important.

DAP-seq binding sites were less important in predicting wound response than the known sequence-based sites, but still ranked among the top 10 most important features for models predicting early wound response (**Table S5**). For example, the CAMTA TF family binding site, AAGCGCGTG, was ranked 3rd most important for genes up-regulated 0.25 or 0.5 hours after wounding but dropped to 11th at 1 hour after wounding, and even lower in later time points. At 1 hour, the AP2EREBP TF family binding site, GGCGGCGGCGG, started to become more important, ranking 10th in the model and increasing to 4th at 3 hours after wounding. In contrast, all DAP-seq sites became less important, or not important at all, for predicting genes up-regulated 6, 12, and 24 hours after wounding. Because known sites in response to wounding were found under wound stress but DAP-seq sites were not, this highlights stress-specific sites are more important than general TF binding sites. These findings also show temporal differences in wounding regulation, but also that in-vitro TF binding sites do not capture the entirety of how wounding response is regulated, especially at later time points.

In addition to known *cis-*regulatory elements and DAP-seq sites, open chromatin sites (DHS) were important for predicting expression at all time points after wounding (top ranked DHS sites for each cluster ranged from rank 1∼4). However, DHS features tended to become more important at later time points (**Table S5**). For example, while different types of features were important at earlier time points, at 24 hours after wounding, the top 12 most important features were all DHS-related. This is rather intriguing and suggests epigenetic modifications may be a more important for later response to wounding. We propose three potential explanations for this finding. The first is that at the late time-point transcription factor binding is no longer the major determinant of regulation since the chromatin state for wounding stress has been established. While chromatin state does change under JA or wound stress, it is not clear to what extent or for how long (Berr et al., 2012). Second, at this later time point, the functional diversity of genes expressed has increased so that their transcriptional regulatory mechanisms have become more complicated and thus no single CRE or DAP-seq feature can be found with high importance. The third possibility is that the known CRE or DAP-seq features important for later time points are not present in our dataset. This could be because the later time points are not as well studied or because DAP-seq data is only available for ∼38% of known TFs (Weirauch et al., 2014; O’Malley et al., 2016), which suggests that novel regulatory sequences that have not yet been identified are important regulators of wound response, especially for later stress response.

### Finding important temporal putative cis-regulatory elements for wound response using machine learning

Although known CREs, *in vitro* TF binding data, and DHS are useful for building wound response prediction models for the clusters in **Figure 2**, the model performance is far from perfect which raises the question whether additional CREs remain to be discovered that can better explain the gene expression patterns seen. To discover novel putative CREs (pCREs), a *k*-mer finding approach was used (modified from Liu et al., 2018), where all possible 6-30-mer sequences were tested for enrichment (*p*<0.01, see **Methods**) in the putative promoters of genes for each cluster (see **Methods**). Based on this criterium, between 42-1,081 pCREs were identified as enriched in genes from each wound response cluster, with the exception of the down-regulated wounding after 12 hours cluster, which had no enriched pCREs (for enrichment of pCREs, see **Table S6**). For each wound response cluster, the pCREs were used to build a wound response prediction model. We found that models built with pCREs alone (e.g. RF algorithm, F-measure range = 0.73-0.81) perform better than models built with known CREs, DAP-seq and DHS for all clusters (e.g. RF algorithm, median F-measure range = 0.66 to 0.74, **Figure 2, Table S4**). Interestingly, models that combined pCREs with known CREs, DAP-seq, and DHS data did not perform better than pCRE-based models alone (e.g. RF-algorithm, median F-measure = 0.67-0.80). This indicates that these pCREs, some are variants of known CREs and other novel, may contribute substantially to the regulation of wound response at different time points.

To understand why the models improve with the addition of pCREs, and what impact pCREs have across wounding time points relative to known information and open chromatin sites, we looked at the relative importance (normalized importance score, see **Methods**) of pCREs, DAP-seq sites, and chromatin accessibility sites across the post-wounding time course (**Figure 3)**. Overall, DHS sites tended to be most important, followed by pCREs, and then finally by DAP-seq sites. When looking closer at individual up-regulated clusters, we found for early time point clusters (0.25, 0.5, and 1 hour after wounding, **Figures 3A-C**), a small percentage of pCREs have higher or as high of an importance score as the most important DHS sites. This indicates that a few unique regulatory elements, or variations of known elements which are not captured by the known TF binding sites, are important for distinguishing differential expression at early time points. Other than this small subset of pCREs, DH-sites are in general more important than the majority of pCREs or DAP-seq sites. For middle range to later time points (3, 6, 12, 24 hours after wounding, **Figures 3D-F**), DH sites have the highest importance, but certain pCREs are also important, ranking just below the most important DH sites. In fact, there is a general shift where the majority of pCREs are of higher importance at these time points than at earlier time points (**Figure 3G**). This indicates that even though the most important features at late time points are the DH sites, there are more putative TF binding sites that distinguish mid-range to late wound induced gene expression compared to earlier time points.

**Figure 3.**
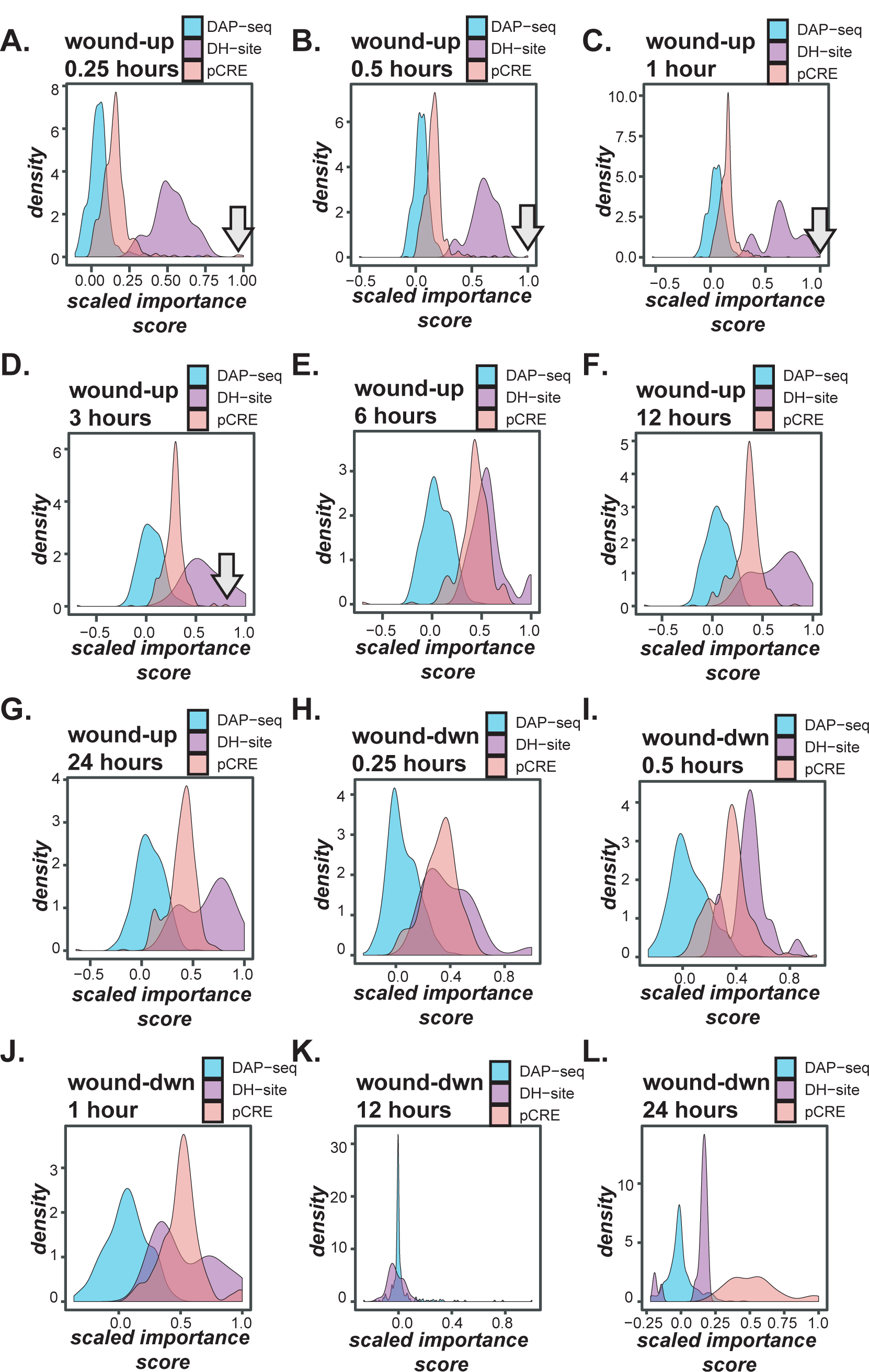
Scaled importance value for each up-regulation wounding time point model (rows) for all features used in the final model. For A-L, density plots show importance value on the x-axis and the density of the feature on the y-axis. DAP-seq sites are in blue, DH sites are in purple, and pCREs are in pink. The importance value is scaled from -1 to 1 for each model, where positive value correlates with differential expression in a given cluster and -1 correlates with the null cluster. The higher absolute value correlates with higher importance of a given feature for a given model. Arrows point to a small peak of pCREs in figures A-D. A. wound up-regulated at 0.25 hours model, B. wound up-regulated at 0.5 hours model, C. wound up-regulated at 1 hour model, D. wound up-regulated at 3 hours model, E. wound up-regulated at 6 hours model, F. wound up-regulated at 12 hours model, G. wound up-regulated at 24 hours model, H. wound down-regulated at 0.25 hours model, I. wound down-regulated at 0.5 hours model, J. wound down-regulated at 1 hour model, K. wound down-regulated at 12 hours model, and L. wound down-regulated at 24 hours model.

Finally, down-regulation importance patterns are less consistent (**Figures 3H-L**). While genes down-regulated at 0.25, 0.5, and 1 hour are fairly similar where either DH sites or pCREs are the most important for regulation, down-regulated genes at 12 hours have no important pCREs, and down-regulated genes at 24 hours have almost exclusively pCREs as being important for regulation. This indicates that 24 hours after wounding is under the most unique regulation compared to all other time points, where up-regulated genes are mostly regulated by open chromatin, and down-regulated by pCREs. Also, it is important to note that open chromatin sites appear to be important features for models at all time points. While some sites are more important to earlier time points than they are to later ones, many sites have an equal relative importance across wounding time point, indicating open chromatin sites cannot distinguish the different expression patterns at different times after wounding. Together, the most distinguishing regulation of wounding time points may be the pCREs.

### Correlation to transcription factor families and cis-regulatory differences across time

Next we wanted to determine which pCRE were similar to a known TF binding motif and which were likely to be novel regulatory elements. To do this, we first calculated the sequence similarity between each pCRE and each known binding motif in order to find the TF family who’s binding motifs best matched the pCRE. **Figure 4** shows the importance rank across all time points for the top 10 most important pCREs for each wounding model. Similar to how more of the same genes were differentially expressed at nearby time point (i.e. the cascade effect), we found more important pCREs were shared with close time points, however some pCREs were uniquely important at a single time point. In fact, none of the top 10 most important pCREs were shared across all time points (for the importance rank of pCRE and PCC to known TF binding motifs from each TF family, see **Table S7)**.

**Figure 4.**
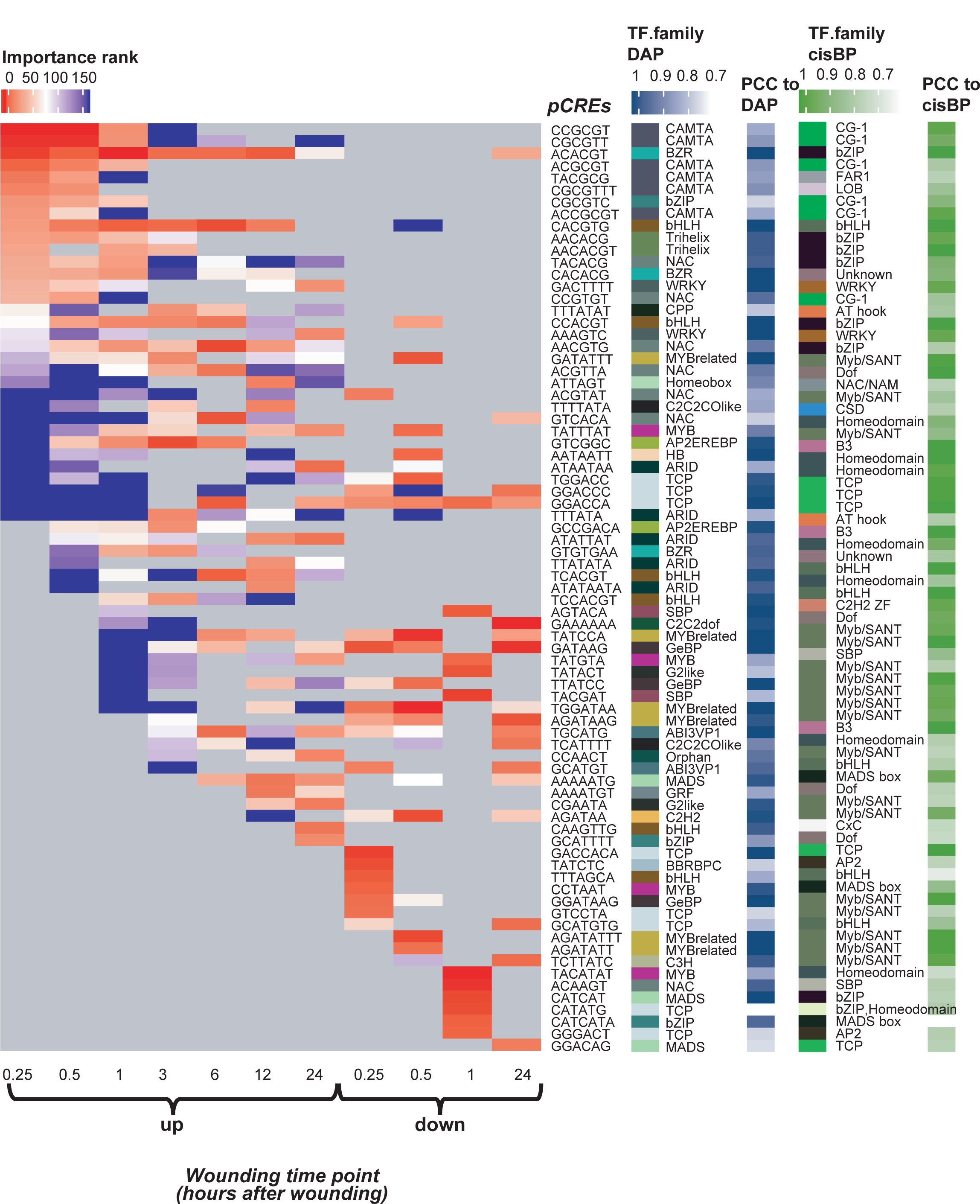
Average importance rank for the top 10 pCREs for each wounding time point model and their TF family. Wound time point models are the columns while pCREs are the rows. Average importance rank is the average rank across five duplicate models ran for the same time point. Highest rank (1) is red and ranks 150 or lower are blue. TF family association is based on the maximum PCC to known TF binding sites. PCC is shown for both DAP-seq and TFBM sites.

For early time points after wounding (i.e. 0.25, 0.5, and 1 hour), many of the top important pCREs were shared and resemble TF binding sites in the *CG-1, bZIP, FAR1, LOB*, and *bHLH* TF families (right two panels; **Figure 4, Table S7**). Different binding sites which bind multiple TF families is consistent with the notion that a variety of signals are induced by wounding. For example, while JA is induced by wounding, other hormones or signals involved, such as ACC, hydrogen peroxide, and ABA, can amplify the JA response (Howe, 2004).

Focusing on the top 3 most important pCREs for each time point (excluding DAP-seq or DH sites, **Figure 5**), we found that 0.25 and 0.5 hours after wounding, CCGCGT, which is most similar to the binding motif of a *CG-1* TF family TF, was the most important pCRE for up-regulated gene models, it then dropped to the 29^th^ most important pCRE at 1 hour after wounding, and by 6 hours after wounding is not enriched in the cluster (**Figure 5, Table S7**). This binding site is associated with TFs which respond to both abiotic and biotic stress. On the other hand, one hour after wounding, the pCRE CACGTG, which was not as important previously (0.25 hour rank = 30, 0.5 hour rank = 17), was the 8^th^ most important pCRE. This pCRE was most similar to the known binding motif for *Myc2*, a *bHLH* TF that responds to both JA and wounding (Dombrecht et al., 2007) (**Figure 5, Table S5**). This element remained important at both 3 and 6 hours after wounding (ranked 10 and 5, respectively), indicating a change in response at 1, 3, and 6 hour time points and that JA hormones have been activated.

**Figure 5.**
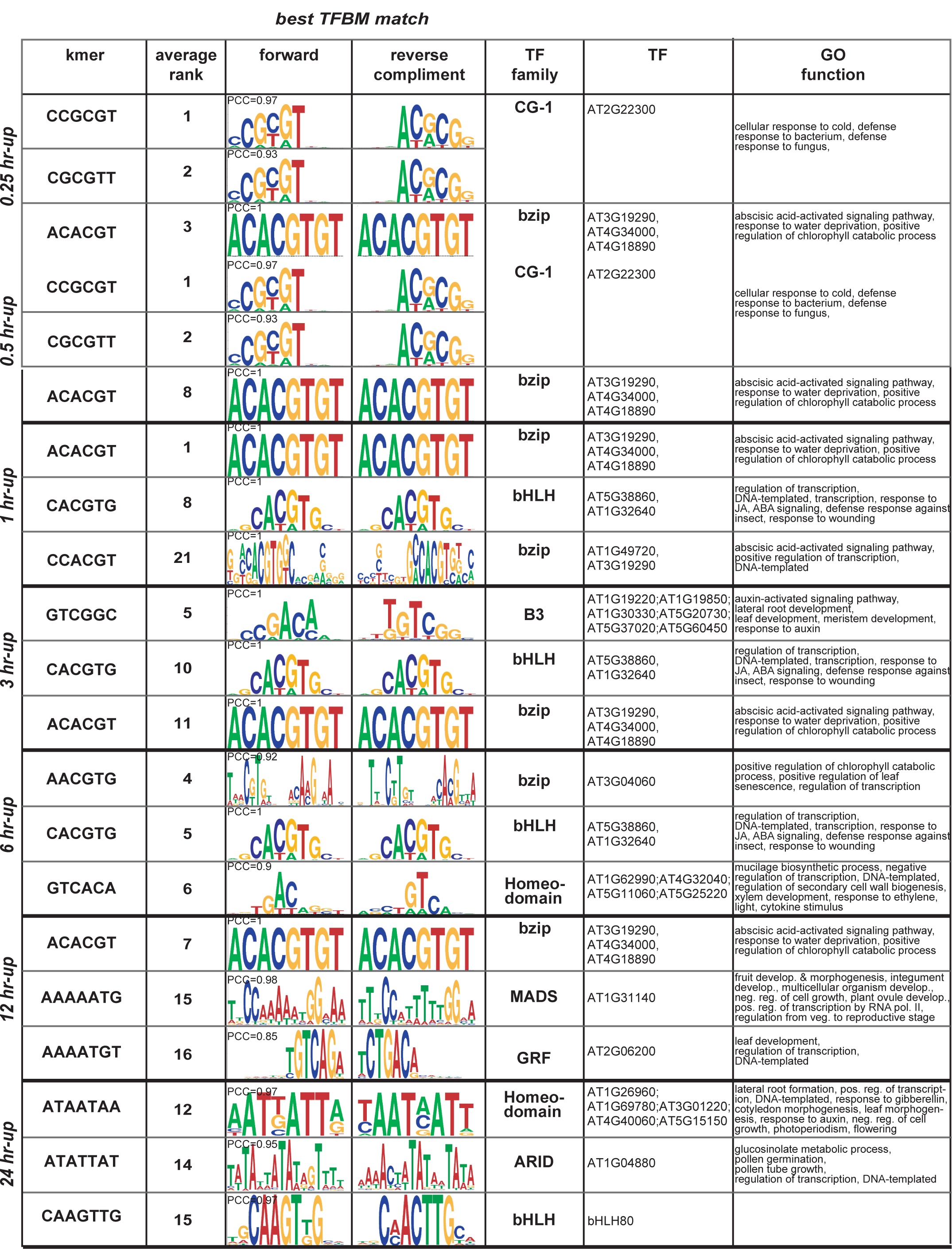
Motif logos for the top 3 pCREs for each up-regulated wounding time point. Chart is divided by time point (0.25 to 24 hours after wounding). The first column is the top 3 ranked pCREs for that time point. The second column is the average rank for that pCRE in the given model. The third and fourth columns are the best matched TF binding motif logos, forward and reverse compliment with PCC value. Columns 5-7 are the TF which binds a given logo (column 6), the TF family the TF belongs to (column 5) and GO functions of the TF (column 7).

Other important early pCREs remained important across the wider range of time points. One example is ACACGT, a pCRE most similar to the known binding motif for *bZIP* family TFs, which are activated by ABA (Yamamoto et al., 2011) and regulate responses to water deprivation (**Figure 5**). This pCRE was enriched in the promoters of genes from all time points and was important (rank < 11) for models of wounding response at 0.25, 0.5, 1, 3, 6, and 12 hours (**Figure 4, Table S7**).

Two pCREs (GTCGGC and GTCACA) were uniquely important for models built for mid-range time points (i.e. 3 and 6 hours after wounding), as the 5^th^ and 6^th^ most important pCREs for the genes up-regulated at 3 hours and 5^th^ and 18^th^ most important at 6 hours (**Figure 5**). These elements were most similar to binding motifs of B3 and Homeodomain family TFs, respectively. Given these TF families are involved in development, response to auxin, and secondary wall biogenesis, this indicates that by 3 to 6 hours after wounding, the damage is likely being repaired.

At the latest time points (i.e. 12 and 24 hours after wounding), we found that some important pCREs were the same as the pCREs important for earlier time points, while others were unique to the later response. As discussed previously, ACACGT, which was ranked 7th at 12 hours after wounding and was also important for earlier time points (0.25, 0.5, 1, 3, and 6 hours after wounding). While ATATTAT, which was most similar to binding motifs of TFs in the ARID family, was ranked 14^th^ at 24 hours after wounding (**Figure 5, Table S7)** This TF family is involved in regulating glucosinolate metabolism. Other important pCREs at the latest time points, ATAATAA and AAAATGT, were elements that bind TF families which regulate development (**Figure 5, Table S7**).

In summary, we found that pCREs important for our models of response at early time points (0.25 to 0.5 after wounding) tend to be associated with many stress and hormone responses, while pCREs 1 to 6 hours after wounding tend to be associated with TFs involved in JA signaling and ABA signaling. Finally, from 3-24 hours after wounding the pCREs tend to be associated with TFs involved in growth and very late responses (12-24 hours after wounding) are associated with TFs related to metabolic defense. Overall, we generated models of the cis-regulatory code in response to wounding that demonstrate how different sets of pCREs, which are likely bound by a variety of TFs, are important at different response times after wounding and could work to regulate a dynamic response to wounding over time.

### Experimental validation of important unknown CREs in early wound response

We validated our model by using the CRISPR/Cas9 system to evaluate the biological significance of the pCRE in *planta*. The top (most important) pCRE found for 0.25 and 0.5 hours, and still important 1 hour after wounding was CCGCGT. Thus, we chose CRISPR/Cas9 target promoters that contain the CCGCGT motif and sorted out the candidates by the following criteria. First, the expression level of the gene, which has the target motif on its promoter region, was relatively high in log fold change at the early time points in response to wounding stress.

Second, the pCRE site is near the PAM sequence such that the site renders susceptible to the CRISPR/Cas9 mutation (Osakabe et al., 2016). We finally selected genes *JAZ2* and *GER5. GER5* was highly expressed and contained the CCGCGT motif. *JAZ2* is a well-known JA-responsive gene (Fernández-Calvo et al., 2011) and the promoter region contained the G-box motif (CACGTG), which can be utilized as a positive control in our mutation assay (Figueroa and Browse, 2012), as well as the CCGCGT motif.

Next, we made the CRISPR/Cas9 construct that targets the pCREs in *JAZ2* and *GER5* promoters and transformed those into Col-0. From antibiotic resistance T_1_ plants, we found a homozygous mutant called *jaz2-4ger5-3/Col-0*. The *jaz2-4ger5-3* double mutant had one base pair insertion in the CCGCGT motif on both *JAZ2* and *GER5* promoters, where T insertion in the *JAZ2* promoter led to no significant nucleotide change from CCGCGT to CCGCGTT, while G insertion in the *GER5* promoter caused a base alteration from CCGCGT to CCGCGGT (**Figure 6**). Further, we generated a homozygous mutant called *jaz2-5/Col-0* that harbored a mutation within the G-box motif of *JAZ2* promoter, in which the CACGTG motif was mutated to CACGTTG (**Figure 6**). To determine the effect of the motif mutations on their downstream gene expression upon wound treatment, we harvested the seedlings of *jaz2-4ger5-3* and *jaz2-5* mutants, as well as Col-0 controls, 1 hour after wounding. The transcript abundances of both mutants and Col-0 were analyzed by reverse transcription-quantitative PCR (RT-qPCR) analysis.

**Figure 6.**
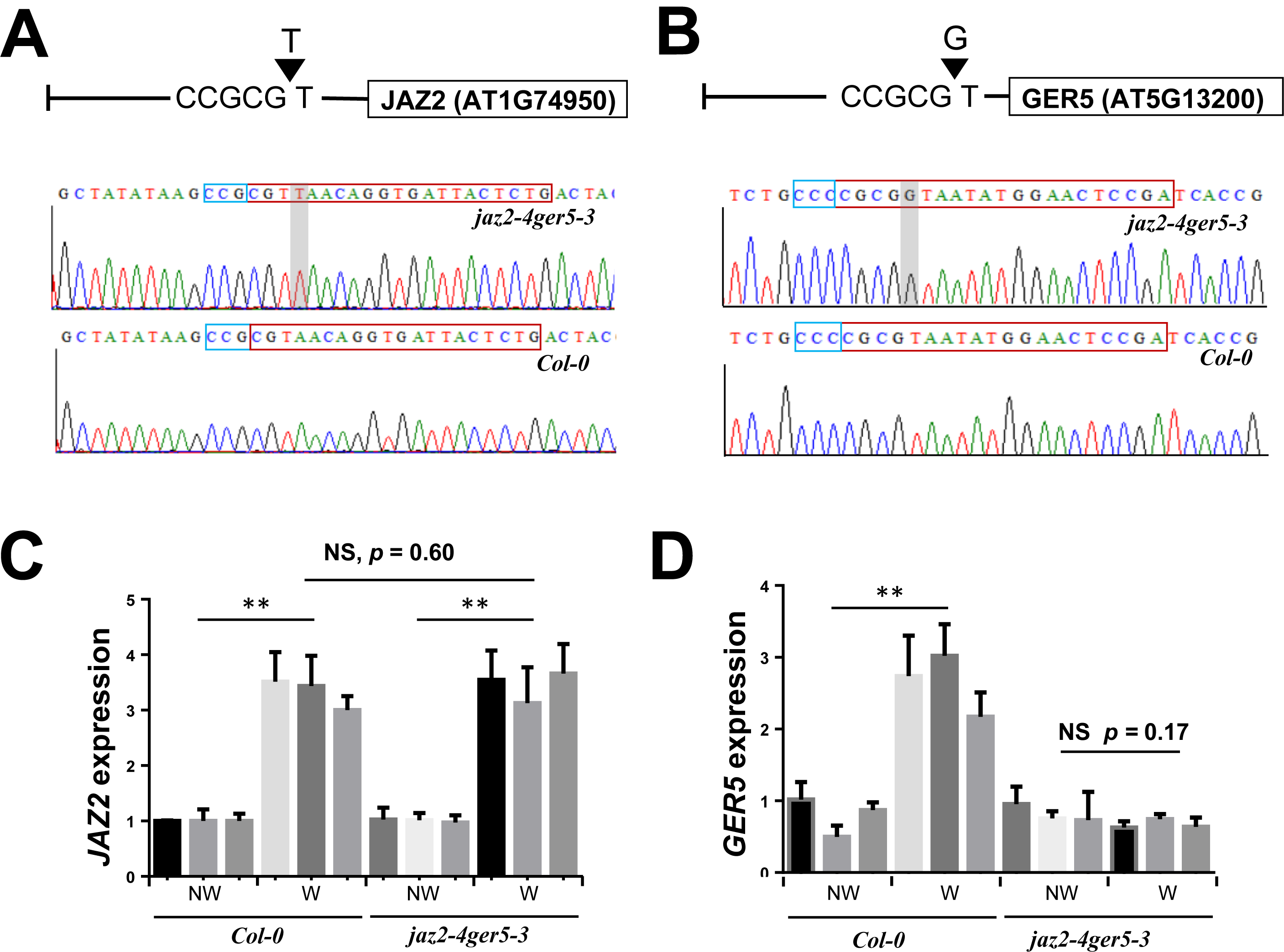
Mutation in predicted *cis*-regulatory motif abolished GER5 induction by wounding. **A**. CRISPR/Cas9 mutation in the CCGCGT motif of in the JAZ2 promoter region (resulting in jaz2-4 ger5-3 line). **B**. CRISPR/Cas9-mediated mutation in the CCGCGT motif of the GER5 promoter in jaz2-4 ger5-3. Chromatogram represent the sequence of the JAZ2 or GER5 promoter regions targeted by CRISPR/Cas9 (upper chromatograms), and corresponding regions in Col-0 (lower chromatograms). Shadowed area denotes the T or G insertion within the CCGCGT motif. Blue and red boxes indicate PAM sequence and gRNA target regions, respectively. **C**. Wound-induced JAZ2 expression in Col-0 and in jaz2-4 ger5-3. **D**. Wound-induced GER5 expression in Col-0 and jaz2-4 ger5-3. Transcript abundances of JAZ2 or GER5 evaluated by RT-qPCR were normalized to ACTIN2. NW and W indicates no wound and wound treatment, respectively. Biological triplicates are shown by individual bars with error bars depicting three technical repeats for each biological replicate. Significant differences levels were analyzed by one-way analysis of variance (ANOVA) and indicated with asterisks [Non-Significant (NS) p > 0.05, * p < 0.05, ** p < 0.01).

We found in the double mutant *jaz2-4ger5-3* the expression of the *JAZ2* gene was up-regulated after wounding, exhibiting the same phenotype as the Col-0 control (**Figure 6**). Thus, as expected, *JAZ2* was not altered in *jaz2-4ger5-3* by wounding, because the CRISPR-Cas9 mutation had resulted in synonymous change. Interestingly, the expression level of *GER5* was not changed in *jaz2-4ger5-3* upon wound treatment, while the expression of *GER5* was significantly up-regulated in the Col-0 control. This indicates that CCGCGT is a novel functional motif that enables the *GER5* gene to respond to early wounding. CCGCGT is in the CAMTA related TF binding family (**Table S7**), which has been shown to respond to cold treatment and may impart freezing tolerance in *Arabidopsis thaliana* (Doherty et al., 2009).

The *JAZ2* expression was up-regulated in response to the treatment in both Col-0 and in the *jaz2-5* mutant. Additionally, the *GER5* transcript level was induced after wounding the in *jaz2-5* mutant (**Figure S2**). In the case of JAZ2, the G-Box CRE CACGTG was changed to CACGTT, which is actually a G-Box variant (Dombrecht et al., 2007). Thus, while significant, the change did not substantially alter the JAZ2 response (effect size=) compared to the *jaz2-4ger5-3* mutant (effect size=). The *GER5* transcript was also not markedly different in the jaz2-5 mutant relative to the Col-0 response (effect size, p-value), indicating the intact CCGCGT CRE regulates *GER5*.

### Modeling the regulatory code of JA-induced and non-JA-induced response to wounding

Having demonstrated how wounding response regulation changes over time and that putative CREs identified as important by the model can impact wound response, we next wanted to study the regulatory differences between JA-induced and non-JA-induced wounding responsive genes. Non-JA-induced wounding responses include those induced by RNase and nuclease activities that are triggered by wounding but not the application of JA (LeBrasseur et al., 2002). Thus, to understand how non-JA induced wound responses are regulated, we used the hormone treatment data described above (Goda et al., 2008) to identify genes that were differentially expressed in response to wounding but not in response to JA at each timepoint. For this analysis, we only included timepoints (30 minutes, 1 hour, and 3 hours) for which we had data for both JA treatment and wounding. Across these three timepoints only 16%, 26%, and 28% of genes up-regulated after wounding were also up-regulated after JA treatment, respectively (**Figure 7A**). The large number of non-JA induced wounding responsive genes is consistent with other studies of wounding which relay a JA-independent mechanism for wounding response (León et al., 1998; LeBrasseur et al., 2002). However, the regulatory code that governs these responses is less well understood.

**Figure 7.**
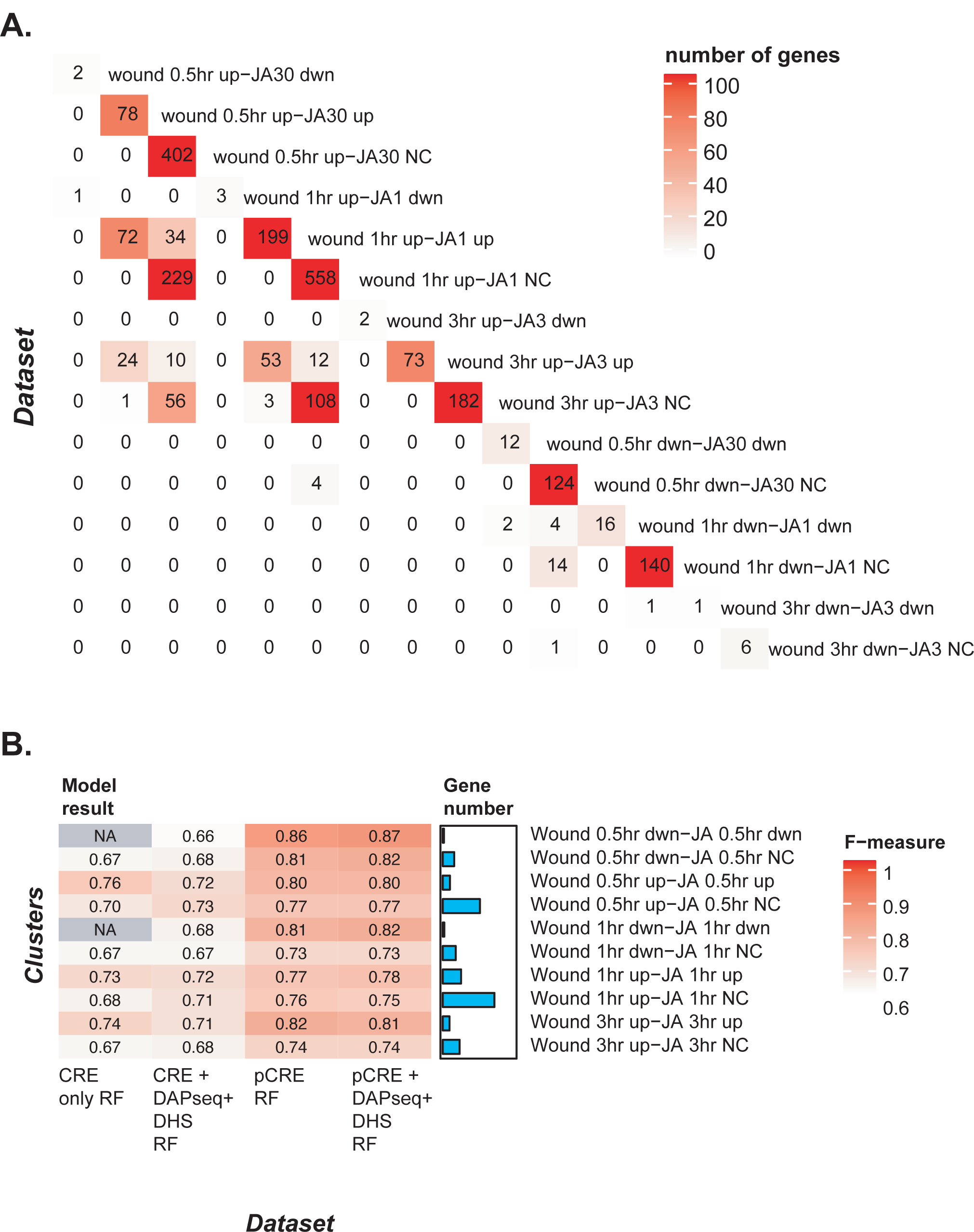
Gene overlap and model performance of each wound JA-induced and wound non JA-induced cluster. A. Heatmap showing the number of genes overlapping in each wounding-JA cluster. The order of rows and columns are the same, based on time point. Number of genes range from 0 (white) to 558 (red) and actual value is printed in the heatmap. B. Each row is a different cluster (JA-induced or JA non-induced) which was used to build a separate model. Each column represents the datasets used as features in the model and the algorithm used (RF= Random Forest). Known only refers to CREs found in the literature (**Table S3**). DAPseq and DHS refer to the DAP-seq and Dnase I hypersensitivity sites. FET enriched 6mer refers to the pCREs which were enriched for a specific cluster. The F-measure range is from 0.5 (white) to 1 (red), and gradient as well as actual F-measure is shown in each cell. The bar chart next to the heat map corresponds to each row/cluster and represents the number of genes in that cluster.

To determine the regulatory differences between JA-induced and non-JA induced wounding response, we first needed to generate models of these different regulatory codes. Using the same approach as described above, we generated machine learning models of wounding response based on known CREs, DAP-seq sites, DH sites, and pCREs for up- and down-regulation responses at different time points. However, here we divided our genes in each cluster further by if they were differentially regulated under wounding and JA treatment (JA-induced) or under wounding but not JA treatment (non-JA induced). Note that models were not generated for genes down-regulated 3 hours after wounding because not enough genes were available for training. Similar to our earlier results, we found that pCRE based models (F-measures: 0.73 ∼ 0.87) performed better than both known CREs based models (0.67 ∼ 0.74) and known CREs and DAP-seq and DHS based models (0.66 ∼ 0.73; for RF models: **Figure 7B**, for SVM models: **Figure S1B, Table S4**). This, again, indicated that pCREs were better able to model the regulation of JA-induced and non-JA-induced wounding response across time points, than using only known TF sites.

### Identifying differences between JA-induced and non-JA-induced CREs: Known and putative

To better understand differences between how JA-induced and non-JA-induced wounding responses are regulated, we next compared the importance of known CREs, DAP-seq sites, DH sites, and pCREs across models. We identified differences in which known CREs were important at 30 minutes and 1 hour after wounding between JA-induced and non-JA-induced responses. For example, CGCGTT, the *RWR* element, was the most important element for the non-JA-induced model, while for JA-induced models, the most important element was the *Myc* element, CACGTG (see **Table S8** for pCREs, DH, and DAP sites and their respective importance scores for non-JA and JA induced models). Interestingly, the *Myc* element also ranks as the third most important feature in the non-JA-induced models. This could be because other TFs not involved in JA response (e.g. *Myc*-LIKE and *BIM3* TFs) can bind to this element (O’Malley et al., 2016) or because the *Myc* element may be necessary to facilitate TF binding to a different regulatory element important for non-JA-induced response. Finally, we found that chromatin accessible sites have a higher overall importance for non-JA-induced than for JA-induced wounding response. Out of the top 10 most important features, 4 to 8 were DH sites for non-JA-induced up-regulated models. In contrast, for JA-induced up-regulated models, none of the top 10 most important features were DH sites (**Table S8**). This indicates that open chromatin sites are important for distinguishing genes that are non-JA-induced from those that are JA-induced.

Next, we compared the importance of pCREs between JA-induced and non-JA-induced models and, with the exception of the G-box motif (CACGTG) and the bZIP binding site (ACGTGT), found there to be little overlap between the two (**Figure S3**). For example, AACGTG and CACGTTT were ranked from 1st to 7th across timepoints in JA-induced models but were not enriched or were ranked much lower (69th to 157th) for non-JA-induced models (**Figure S3, Table S8**). These pCREs were most similar to binding motifs of TFs in the NAC and CAMTA families, respectively. In contrast, CCGCGT and GCCGAC, were the most important pCREs 0.5 and 3 hours after wounding in the non-JA-induced models but were not enriched or were ranked much lower (232th importance) for JA-induced models **(Table S8**).

These pCREs were most similar to the binding motifs of TFs in the CG-1 and B3 TF families, respectively. Interestingly, these TF families are known involve TFs that have a general response to stress as well as those which are involved in auxin signaling and development, indicating two functions of wound stress response which do not involve JA. Together, this highlights how JA-induced and non-JA-induced differential gene expression is likely regulated by different sets of regulatory elements that are recognized by different families of TFs.

### Modeling SM pathway regulation using wound stress data

Another way to study response to wounding is by focusing on the response of whole metabolic pathways instead of the response of individual genes. Here, we measured the degree to which genes annotated as belonging to a particular specialized metabolism pathway were enriched in the genes up-regulated across the time series (**Table S9**). At earlier to mid-range time points (from 0.25 to 3 hours after wounding) JA biosynthesis was the most enriched pathway (*p*-values range from 0.0015 to 3.5e-07; **Table S9**). However, by 6 hours after wounding, JA biosynthesis genes were less enriched (*p*-values = 0.0018) and by 12 hours it is not enriched at all. This demonstrates how the JA biosynthesis pathway is only activated early in wounding response. Genes from the glucosinolate biosynthesis from tryptophan (Gluc-Trp) pathway, on the other hand, were enriched 0.5 hours after wounding (*p*-value = 0.008), were most enriched 12 hours after wounding (*p*-value = 0.0008) and were not enriched by 24 hours after wounding (*p*-value = 0.1). Finally, two genes from the anthocyanin biosynthesis pathway (AT2G38240 and AT5G05600) were up-regulated 0.5 hours after wounding and the same two genes remain up-regulated through 24 hours after wounding. These examples demonstrate that some wounding responsive pathways are dynamic over time, while other wounding responsive pathways are steady.

To determine how dynamic changes in a metabolic pathway are regulated, we used the glucosinolate biosynthesis from tryptophan (Gluc-Trp) pathway as an example. No Gluc-Trp pathway genes were up-regulated at the earliest time point, however by 0.5 hours after wounding three genes were significantly up-regulated and by the one hour time point three additional genes were significantly up-regulated (see stars; **Figure 8A**). Looking beyond the first hour, we saw a cascading effect, where by 3 hours after wounding, the genes turned on at one hour still were still up-regulated, but the three genes that were first up-regulated at 0.5 hours were turned off. Continuing this trend, by six hours after wounding, only one gene that was up-regulated at one and three hours after wounding was still significantly up-regulated. (**Figure 8A**). This type of pattern could be due to genes upstream in the pathway being involved in up-regulating genes downstream in the pathway or could be due to having different TFs, not in the Gluc-Trp pathway, regulating up and downstream pathway genes.

**Figure 8.**
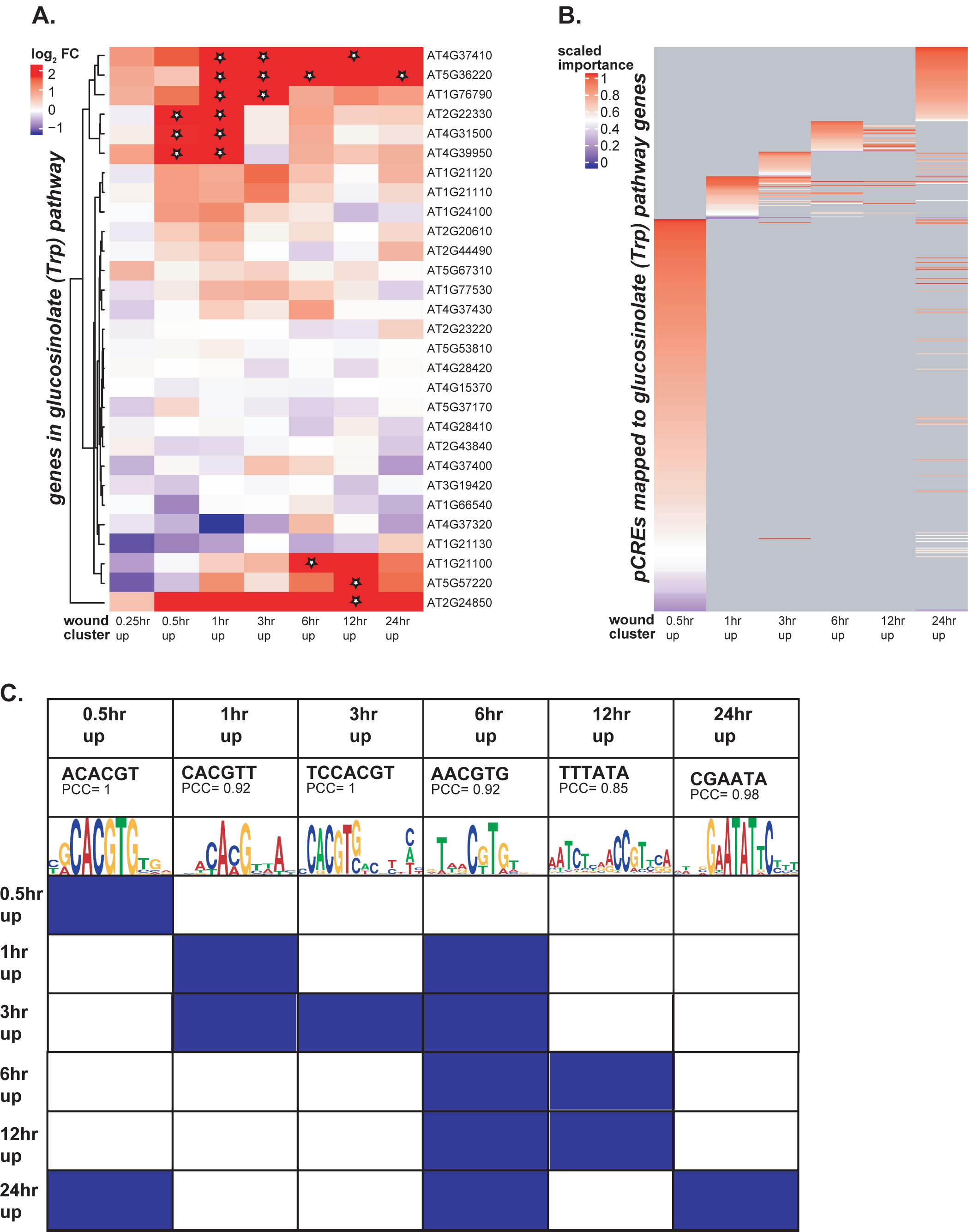
Co-expression and regulation of Glucosinolate from Tryptophan pathway genes. A. Heatmap showing the log_2_ fold change values of all genes in the Gluc-Trp pathway across the 7 wounding time points. Genes are clustered using hierarchal clustering. Genes are on the y-axis, wounding time points are on the x-axis, and log_2_ fold change is represented as the color gradient from a value of 2 or greater (red) to a value of -1 or less (blue). Stars indicate gene is significantly up-regulated at a given time point. B. Scaled importance score of pCREs mapped to Gluc-Trp genes which are up-regulated at a given wounding time point. Importance is scaled from 0 to 1, where 1 is most important and 0 is least important. Each row is a pCRE and each column is the wounding time point. C. The most important pCRE for Gluc-Trp pathway genes at a given time point. First row is the pCRE and correlation to a known TF binding site. The second row is the motif logo for the known TF binding site. The rest of the rows show whether that particular pCRE overlaps with Gluc-Trp genes at other time points.

To understand how the cascading response is regulated, we mapped the pCREs found from each of the models for up-regulated genes at a given time point back to the promoters of the Gluc-Trp pathway genes (see Methods). **Figure 8B** shows the overlap of pCREs discovered from the wounding time point models that map to Gluc-Trp genes and their importance level at a given time point. We can see that starting at 0.5 hours after wounding, there is little overlap of important pCREs across time points with the exception of pCREs present at 6 and 12 hours after wounding. This indicates that for the Gluc-Trp pathway, genes turned on at different times have different regulatory elements which are specific to those genes. **Figure 8C** shows the highest ranked important pCREs for Gluc-Trp pathway genes at a given time point, and whether this pCRE is present or absent in Gluc-Trp pathway genes at other time points. For example, ACACGT, which is perfectly similar to the binding motif of a TF from the *bZIP* family (PCC=1), when specifically finding pCREs in Gluc-Trp genes, this pCRE is the most important element at 0.5 hours after wounding (**Figure 8C**). Additionally, this element is not found in Gluc-Trp pathway genes up-regulated at other time points. Other pCREs are important for regulating expression of pathway genes at later time points. For example, AACGTG, which is most similar to the binding motif of a *bZIP* family TF, is enriched in the promoters of Gluc-Trp pathway genes up-regulated 1, 3, 6, 12, and 24 hours after wounding, but has the highest importance at 6 hours after wounding. Overall genes belonging to the Gluc-Trp pathway have varied cis-regulatory elements depending on when they are up-regulated after wounding, and timing of response can be an important consideration when finding CREs related to certain pathways.

## Conclusion

The aim of this study was to better understand the temporal differences in transcriptional response to wounding stress in *A. thaliana*. We accomplished this by integrating multiple levels of regulatory information (e.g. sequence based and epigenetic features) into machine learning models of the regulatory code that could be used to predict if a gene was up- or down-regulated at a specific timepoint after wounding. We demonstrated that wounding response is regulated by a diverse set of regulatory elements that are likely bound by TFs from a wide range of TF families. We identify 4,255 pCREs derived from wounding co-expression clusters up-regulated at different timepoints, with 3,493 (82%) having significant sequence similarity (PCC > 0.8) to known TF binding sites. These pCREs were more predictive of differential expression at each wounding time point than models based on known TF binding sites (derived from the literature and the DAP-seq database) and information about open chromatin sites. From our machine learning models, we were also able to quantify the relative importance of each pCRE included in the model for each time point. While some pCREs were important across multiple timepoints, we generally found that pCREs were either important for early or late time points after wounding.

By modeling JA-induced and non-JA-induced transcriptional responses separately, we were able to identify 2,569 pCREs important for predicting genes up-regulated in response to wounding but not in response to JA treatment. Of these, 2,371 (92%) had significant sequence similarity (PCC > 0.8) to known TF binding sites. In addition, by focusing on genes in the Gluc-Trp pathway, we were able to identify pCREs important for predicting genes in this wound responsive specialized metabolite pathway.

While our models perform notably better than random expectation, there is room for improvement. One possible reason we could not predict differential expression perfectly is that we limited our study to focus on CRE sites in the promoter region (+1kb upstream of the transcription start site). However, CREs can be located in other regions, including in the downstream untranslated regions of the gene, in introns, or coding regions (Rose et al., 2008), which could be evaluated in future studies. Another limitation is that genes up- or down-regulated at a particular time point might not all be regulated the same way. This is especially likely for large time point gene groups, like the cluster of up-regulated genes one hour after wounding, which contains 760 genes. If we could further break down this group, perhaps based on the gene’s response to other stresses, we may be able to model more specific responses at one hour, which could improve the overall performance. Finally, data regarding DAP-seq and DH sites did not come from wounded plants, and therefore are not capturing any changes that may occur to chromatin state or TF-binding sites after wounding.

Many of the important pCREs found in this study have not been shown to be associated with wounding. This is especially true for pCRE found at later timepoints that have been less well studied. However, new technologies, such as CRISPR-cas9, make it possible to generate precise edits to the DNA allowing for the role of these pCREs in temporal wounding response to be tested experimentally. We were able to successfully mutate the pCRE CCGCGT and this resulted in a drop in expression under wounding in the target gene GER5. Thus, one predicted previously untested CRE has now been shown to be involved in wound response. To that end, our study provides a set of important putative targets that could be used to prioritize experiments to can confirm novel pCREs associated with different types of wounding response. Finally, more can be done to find regulatory elements associated with different pathways. Because the Gluc-Trp pathway was associated with wounding, we were able to find elements which may help regulate that pathway. However, other pathways may respond to different types of stress or may be active during certain stages of development or in particular tissues. Therefore, future studies should focus on determining regulatory elements for particular pathways by using an associated expression data set.

## Methods

### Expression datasets and analysis

Microarray data from three different AtGenExpress studies were downloaded from TAIR and CEL files were processed using Affy program in R. The studies included biotic stress (Wilson et al., 2012), abiotic stress (Kilian et al., 2007; Wilson et al., 2012), and hormone treatment (Goda et al., 2008), where wounding is part of the abiotic stress dataset. These studies grew plants under similar conditions, were treated 18 days after germination, and were all part of the AtGenExpress project. Each study had 8 different treatments of either different stresses or hormones, for a total of 24 data sets. Samples from each data set were collected after treatment at a range of time points, including 15 minutes, 30 minutes, 1 hour, 2 hours, 3 hours, 4 hours, 6 hours, 12 hours, and 24 hours after treatment. Note that not all time points were used for each treatment. For each data set, controls were collected at the same time in order to control for circadian effects.

Differential expression was calculated using affy and limma packages in R (Gautier et al., 2004; Ritchie et al., 2015), and significantly differentially expressed genes were those that had an absolute log2 fold change ≥ 1 and adjusted p-value < 0.05. Up-regulated genes were those genes which were differentially expressed but with a log2 fold change ≥ 1, while down-regulated genes were those genes which were differentially expressed with a log2 fold change ≤ -1. For each expression dataset, Pearson’s Correlation Coefficient (PCC) was calculated between each treatment.

### Gene clusters

Wounding time point clusters were determined by differential expression at each time point of wounding stress (0.25, 0.5, 1, 3, 6, 12, and 24 hours after wounding). For example, genes which were up-regulated at the time point of 1 hour after wounding were placed in cluster 1 while genes down-regulated at 1 hour after wounding were placed in cluster 2. This created a total of 14 wounding clusters. For wounding and JA clusters, genes were placed in a cluster based on whether they were differentially expressed in one or both treatments at the same time point. For example, a gene X up-regulated in both 1 hour after wounding and 1 hour after JA treatment would be placed in cluster 1, while gene Y up-regulated in 1 hour after wounding but not changed under 1 hour after JA treatment would be placed in cluster 2. Finally, for a gene Z up-regulated in 1 hour after wounding but down-regulated in 1 hour after JA treatment would be placed in cluster 3. Therefore, at each time point which is in both wounding and JA treatment datasets (0.5, 1, and 3 hours) each up- or down-regulated after wounding cluster was divided into 3 separate clusters, for a total of 18 clusters. Three of these potential clusters actually contained no genes and were subsequently omitted (up-regulated after wounding but down-regulated after JA treatment at 0.5 hours, 1 hour, and 3 hours). A non-differentially expressed cluster was determined by genes which were not differentially expressed across all stresses and timepoints as well as all hormone treatments. For all gene clusters and overlap of clusters, see **Table S1**.

### Known cis-regulatory elements literature search

Known regulatory elements were curated from a literature search. They included elements shown to be responsive to JA, wounding, or insect stress. The studies for this search can be found in **Table S3**. Both experimental and computational data was included.

### Putative Cis-regulatory finding

Promoter regions of each gene (identified as 1-kb upstream of the transcription start site) were downloaded from TAIR for *A. thaliana*. Using homemade python scripts (https://github.com/ShiuLab/MotifDiscovery) were used to identify all combinations of 6-mers present in gene promoters. The Fisher’s Exact Test (FET) was then used to determine overrepresented putative cis-regulatory elements (pCREs) in the promoter region (defined as 1000 bp upstream of gene start site) by comparing a given wounding up- or down-regulated cluster to the non-differentially expressed cluster. A range of p-value cutoffs (adjusted P < 0.01, P<0.01, adjusted P < 0.05, and P < 0.05) was used, however for later machine-learning models, the best results were with the non-adjusted P < 0.01. Using the Motif Discovery pipeline, kmers (oligomer sequences of length k) were searched for in the promoters of genes of interest. Starting with all possible 6-mers, sequences which were found to be significantly overrepresented in the clusters based on the p-values listed above, were kept. Another round of kmer finding then occurred where the significant 6mer was extended on either side, producing two 7mers, and these 7mers were again tested to see if they were significantly overrepresented in the given cluster, and if their p-value was lower than the parent 6mer. If this was true, the 7mer was kept and the 6mer discarded. If not, the 7mer was discarded and the 6mer was kept. This procedure of “growing” kmers continued until the longest kmer with a p-value lower than its predecessor was obtained. These pCREs were then used as features to predict expression in machine-learning models.

TAMO/1.0 (Gordon et al., 2005) was also used to create tamo files for each motif, which was used later to correlate to known transcription factor binding sites.

### Arabidopsis cistrome and epicistrome

Two datasets providing *in-vitro* transcription factor (TF) binding sites were used to correlate to pCREs. First, *A. thaliana* motifs (position weight matrices) determined from protein binding arrays (called TF binding motifs or TFBMs) (Weirauch et al., 2014) were downloaded from http://cisbp.ccbr.utoronto.ca. DNA affinity purification sequencing (DAP-seq) peaks (O’Malley et al., 2016) were downloaded from http://neomorph.salk.edu/PlantCistromeDB. The peaks were then mapped to *A. thaliana* genome using python scripts. If the peak overlapped with the promoter of a gene of interest, the peak was considered present as a feature for that gene. To provide insight into chromatin structure, Dnase I hypersensitivity (DH) sites (Zhang et al., 2012) were obtained from the National Center for Biotechnology Information database under the ID number GSE34318 as bed files. Bed files were parsed using python scripts to obtain gff files, which were then mapped to the *A. thaliana* genome. If the peak overlapped with the promoter of a gene of interest, the peak was considered present as a feature for that gene.

### Machine learning models

Prediction models were built for each wounding time point cluster as well as for wounding-JA cluster where enriched pCREs from the promoter analysis were used as features to predict expression patterns in each expression class (up- or down-regulated genes in each cluster). Random Forest (RF), Support Vector Machine (SVM), and Gradient Boosting, (GB) were the machine learning algorithms implemented for each cluster using Python package sci-kit learn (Pedregosa et al., 2011). Python scripts used to run the models can be found here: https://github.com/ShiuLab/ML-Pipeline. For each model, 10% of the data was withheld from training as an independent, testing set. Because the dataset was unbalanced (ie. 6,855 null genes, 760 up-regulated genes under wounding at 1 hour), 100 balanced datasets were created from random draws of the null gene cluster to match with the number of genes in the differentially expressed cluster. Using the training data, grid searches over the parameter space of RF and SVM were performed. The optimal hyperparameters identified from the search were used to conduct a 10-fold cross-validation run (90% of the training dataset used to build the model, the remaining 10% used for validation) for each of the 100 balanced datasets. Model performance was evaluated using F-measure, the harmonic mean of precision and recall, where precision is defined as the number of true positives divided by the sum of true and false positives, and recall is defined as the number of true positives divided by the sum of true positives and false negatives. Thus, in a binary model, a perfect prediction has an F-measure of 1 and the random expectation is 0.5. The models also have an importance score for each input feature, which is determined by the decrease in impurity of a node in a decision tree, and then averaged across the trees in the forest. Thus, the higher the number, the more important the feature (Breiman, 2001; Louppe, 2014). Importance value was scaled by normalizing based on the minimum and maximum values (where the minimum importance value is subtracted from a given value i, then divided by the difference between the maximum and minimum importance value). Importance rank was taken by ranking taking the importance value and ranking from highest to lowest. Percentile rank is taken by taking the rank of the feature and dividing it by the total number of features.

For each cluster, models with only known (derived from literature) CREs were built (model set 1), then models with known CREs plus DAP-seq and DH site information were built (model set 2). Finally, models with DAP-seq, DH site and enriched pCRE information were built (model set 3). Additionally, for model set 3, five separate models for each wounding time point cluster were run to determine the average importance score for each feature. This was then used to rank the features (pCREs) from most important to least important based on the average of the importance rank for each feature from the five models. Before ranking, reverse compliment pCREs were removed, so that essentially the same pCRE was not ranked twice. To assess random expectation, gene clusters chosen randomly from the expression data sets were enriched for pCREs. These were then used to build machine learning models using the methods above. Random gene clusters were made for genes at n= 30, 50, 100, 150, 200, and 250 at 20 repetitions each. Model results are reported in Table S4.

### Sequence similarity of pCREs to known TF binding sites

To compare pCREs to potential know TF binding sites, pairwise PCC (Pearson’s correlation coefficient) distance between pCREs and TF binding sites (both DAP-seq and TFBMs) was generated using the TAMO program (Gordon et al., 2005). After calculating the PCC distance to all possible TF binding sites, the lowest distance (highest PCC) was determined for each pCRE as its best match. The best match was then used for visualization of the binding site logo.

### Experimental validation using CRISPR-Cas9

To validate our pCRE predictions, we first chose two regulatory elements to validate. The first was the pCRE CCGCGT, which was the most important pCRE in early wound response (1 at 0.25 and 0.5 hours, and 3 at 1 hour after wounding, **Table S6**) as well as a known wound response element, CACGTG, which was also highly ranked in early wound response, although not as high as CCGCGT (**Table S6**). We mapped these CREs to the promoters of wound responsive genes and found of these genes to be of interest: AT1G74950 and AT5G13200.

AT1G74950 is the well-known transcription factor, JAZ2, that responds to wounding and has the known wound response element, CACGTG, as well as the pCRE, CCGCGT in its promoter. AT5G13200, or GER5, is not known to be involved in wounding, but gene does increase expression under early wounding stress and contains the pCRE CCGCGT. Both genes had CRISPR-Cas9 GG sites in their respective promoters close to the CREs in question, facilitating the use of CRISPR-Cas9 as a way to change these elements *in-vivo* as a way to test their functionality.

### Plant growth condition and wound treatment

The *Col-0* and *CRISPR/Cas9* mutants generated in this study were grown in 16:8 photoperiod (Light: Dark) at 23°C. Wound treatments were done when plants were 13 days old XX as in (Kilian et al., 2007). Plants were wounded as in (Koo et al., 2009) and plant tissue was harvested one hour after wounding, frozen in liquid nitrogen, and then stored at -80C until RNA was extracted.

### Plasmid construction

For the construction of CRISPR plasmids, two gRNAs were simultaneously assembled into the pHEE401E vector by the Golden Gate assembly method (Wang et al., 2015). The gRNA sequences used in this study are shown in **Table S10**.

### Validation of CRISPR/Cas9-induced mutation

The CRISPR plasmids were transformed into GV3101 *Agrobacterium* strain, followed by floral dipping into Arabidopsis (*Col-0*). The T_1_ transgenic plants were grown in MS media containing hygromycin (25 mg/L) for three weeks. Genomic DNA was extracted from the rosette leaf of the hygromycin-resistant T_1_ plants, and the promoter regions of *JAZ2* and *GER5* were amplified by genomic PCR using the primers, which are listed in **Table S10**. The sequences of the regions targeted by CRISPR/Cas9 were validated by Sanger sequencing

### RNA extraction and reverse transcription-quantitative PCR analysis

Expression levels of *JAZ2* and *GER5* were analyzed in the T_2_ generation. Total RNA from CRISPR/Cas9 mutants was extracted with RNeasy® Plant Mini Kit (Qiagen, US) following manufacturer’s instruction. Approximately 500 ng of RNA was used for cDNA synthesis with SuperScript™ II Reverse Transcriptase (Invitrogen, US). The transcript levels of *JAZ2* and *GER5* were determined by quantitative real-time PCR (Quantstudio 3 Real-Time PCR, Thermo Scientific, US) using SYBR Green PCR Master Mix followed by manufacturer’s instruction (ThermoFisher Scientific, CA, USA). The C_t_ values of the genes were normalized to those of *ACTIN2*. The PCR primer sets were described in **Table S10**.

### Pathway enrichment and pCRE mapping

Pathway annotations were downloaded from the Plant Metabolic Network Database (https://www.plantcyc.org/). Enrichment tests were performed by using python scripts (https://github.com/ShiuLab/GO-term-enrichment) and the python fisher 0.1.9 package which implements the Fisher Exact test. In order to map pCREs back to the genes in the glucosinolate from tryptophan (Gluc-Trp) pathway, gff files were created which contained the coordinates of pCREs in the promotors of all *A. thaliana* genes. Genes which were part of the Gluc-Trp pathway which were expressed at a wounding time point were matched up with pCREs which mapped to them. Finally, the importance for pCREs which map to Gluc-Trp genes was determined for each wounding time point from the previous wounding models.

## Figure Legends

**Supplemental Figure 1. Heatmap of the F-measure for all wounding SVM models**. Each row is a different cluster which was used to build a separate model. Each column represents the datasets used as features in the model and the algorithm used (SVM= Support Vector Machine). Known only refers to CREs found in the literature (**Table S3**). DAPseq and DHS refer to the DAP-seq and Dnase I hypersensitivity sites. FET enriched 6mer refers to the pCREs which were enriched for a specific cluster. The F-measure range is from 0.5 (white) to 1 (red), and gradient as well as actual F-measure is shown in each cell. The bar chart next to the heat map corresponds to each row/cluster and represents the number of genes in that cluster. A. Wounding time point models. B. wounding JA-induced and non JA-induced models.

**Supplemental Figure 2. Mutation in CACGTG motif of JAZ2 promoter led to down-regulation of JAZ2 expression following wound treatment**.

**A**. CRISPR/Cas9-mediated mutation in CACGTG motif of JAZ2 promoter region in jaz2-5 mutant. **B**. No mutation in CCGCGT motif of GER5 promoter in the jaz2-5. Note that the CRISPR/Cas9 against the GER5 promoter did not result in nucleotide changes in the region. Upper chromatogram shows the CRISPR/Cas9-targeted promoter regions of the JAZ2 and GER5 promoter. Lower chromatogram displays its corresponding region in Col-0. The T insertion within CACGTG motif of the JAZ2 promoter was marked with shadow on the chromatogram. Blue and red boxes highlight PAM sequence and gRNA target region, respectively. **C**. Wound-induced JAZ2 expression in Col-0 and jaz2-5. **D**. Wound-induced GER5 expression in Col- 0 and jaz2-5. The transcript abundances of JAZ2 or GER5 were normalized to ACTIN2. NW indicates no wound treatment and W indicates wound treatment. Three biological replicates are individually demonstrated with error bars, obtained from three technical repeats. One-way analysis of variance (ANOVA) was applied to analyze statistical significances of the gene expression levels, and the differences are indicated by asterisks. (* p < 0.05, ** p < 0.01)

**Supplemental Figure 3. Average importance rank for the top 10 pCREs for each wounding JA-induced and non JA-induced model and their TF family**. Wound time point models are the columns while pCREs are the rows. Average importance rank is the average rank across five duplicate models ran for the same time point. Highest rank (1) is red and ranks 150 or lower are blue. TF family association is based on the maximum PCC to known TF binding sites. PCC is shown for both DAP-seq and TFBM sites.

## Supplemental Data

**Table S1:** Between sample PCC results

**Table S2:** Sample cluster overlap and genes in each cluster

**Table S3:** Known cis-regulatory elements derived from literature

**Table S4:** All machine learning model results

**Table S5:** Feature importance for models using only known elements or sites

**Table S6:** All pCREs enriched for each wounding time point cluster and their p-values

**Table S7:** Summary table for the importance rank of each pCRE for each cluster and their correlation to DAP-seq or TFBM sites

**Table S8:** Overall feature importance score for wounding JA-induced and non JA-induced clusters

**Table S9:** Pathway enrichment for each wounding time point cluster and their p-values

**Table S10:** Primers used for CRISPR-cas9 and qPCR and promoter sequences of experimental genes

## Notes

### Competing Interest Statement

The authors have declared no competing interest.

## References

Asensi-Fabado M-A, Amtmann A, Perrella G (2017) Plant responses to abiotic stress: The chromatin context of transcriptional regulation. Biochim Biophys Acta BBA - Gene Regul Mech 1860: 106–122

Berr A, Ménard R, Heitz T, Shen W-H (2012) Chromatin modification and remodelling: a regulatory landscape for the control of Arabidopsis defence responses upon pathogen attack: Chromatin regulation of plant defence. Cell Microbiol 14: 829–839

Bostock RM, Pye MF, Roubtsova TV (2014) Predisposition in Plant Disease: Exploiting the Nexus in Abiotic and Biotic Stress Perception and Response. Annu Rev Phytopathol 52: 517–549

Breiman L (2001) Random Forests. Mach Learn 45: 5–32

Chung HS, Koo AJK, Gao X, Jayanty S, Thines B, Jones AD, Howe GA (2008) Regulation and Function of Arabidopsis *JASMONATE ZIM* -Domain Genes in Response to Wounding and Herbivory. Plant Physiol 146: 952–964

Dombrecht B, Xue GP, Sprague SJ, Kirkegaard JA, Ross JJ, Reid JB, Fitt GP, Sewelam N, Schenk PM, Manners JM, et al (2007) MYC2 Differentially Modulates Diverse Jasmonate-Dependent Functions in *Arabidopsis*. Plant Cell 19: 2225–2245

Fernández-Calvo P, Chini A, Fernández-Barbero G, Chico J-M, Gimenez-Ibanez S, Geerinck J, Eeckhout D, Schweizer F, Godoy M, Franco-Zorrilla JM, et al (2011) The *Arabidopsis* bHLH Transcription Factors MYC3 and MYC4 Are Targets of JAZ Repressors and Act Additively with MYC2 in the Activation of Jasmonate Responses. Plant Cell 23: 701–715

Frerigmann H, Gigolashvili T (2014) Update on the role of R2R3-MYBs in the regulation of glucosinolates upon sulfur deficiency. Front Plant Sci. doi: 10.3389/fpls.2014.00626

Gautier L, Cope L, Bolstad BM, Irizarry RA (2004) affy--analysis of Affymetrix GeneChip data at the probe level. Bioinformatics 20: 307–315

Glisovic T, Bachorik JL, Yong J, Dreyfuss G (2008) RNA-binding proteins and post-transcriptional gene regulation. FEBS Lett 582: 1977–1986

Goda H, Sasaki E, Akiyama K, Maruyama-Nakashita A, Nakabayashi K, Li W, Ogawa M, Yamauchi Y, Preston J, Aoki K, et al (2008) The AtGenExpress hormone and chemical treatment data set: experimental design, data evaluation, model data analysis and data access. Plant J 55: 526–542

Gordon DB, Nekludova L, McCallum S, Fraenkel E (2005) TAMO: a flexible, object-oriented framework for analyzing transcriptional regulation using DNA-sequence motifs. Bioinformatics 21: 3164–3165

Howe GA (2004) Jasmonates as Signals in the Wound Response. J Plant Growth Regul 23: 223–237

Howe GA, Jander G (2008) Plant Immunity to Insect Herbivores. Annu Rev Plant Biol 59: 41–66

Hutvagner G, Simard MJ (2008) Argonaute proteins: key players in RNA silencing. Nat Rev Mol Cell Biol 9: 22–32

Ikeuchi M, Iwase A, Rymen B, Lambolez A, Kojima M, Takebayashi Y, Heyman J, Watanabe S, Seo M, De Veylder L, et al (2017) Wounding Triggers Callus Formation via Dynamic Hormonal and Transcriptional Changes. Plant Physiol 175: 1158–1174

Kilian J, Whitehead D, Horak J, Wanke D, Weinl S, Batistic O, D’Angelo C, Bornberg-Bauer E, Kudla J, Harter K (2007) The AtGenExpress global stress expression data set: protocols, evaluation and model data analysis of UV-B light, drought and cold stress responses: AtGenExpress global abiotic stress data set. Plant J 50: 347–363

LeBrasseur ND, MacIntosh GC, Perez-Amador MA, Saitoh M, Green PJ (2002) Local and systemic wound-induction of RNase and nuclease activities in Arabidopsis: RNS1 as a marker for a JA-independent systemic signaling pathway. Plant J 29: 393–403

León J, Rojo E, Sánchez-Serrano JJ (2001) Wound signalling in plants. J Exp Bot 52: 1–9

León J, Rojo E, Titarenko E, Sánchez-Serrano JJ (1998) Jasmonic acid-dependent and - independent wound signal transduction pathways are differentially regulated by Ca 2+ /calmodulin in Arabidopsis thaliana. Mol Gen Genet MGG 258: 412–419

Liu M-J, Sugimoto K, Uygun S, Panchy N, Campbell MS, Yandell M, Howe GA, Shiu S-H (2018) Regulatory Divergence in Wound-Responsive Gene Expression between Domesticated and Wild Tomato. Plant Cell 30: 1445–1460

Lorenzo O, Piqueras R, Sánchez-Serrano JJ, Solano R (2003) ETHYLENE RESPONSE FACTOR1 Integrates Signals from Ethylene and Jasmonate Pathways in Plant Defense. Plant Cell 15: 165–178

Louppe G (2014) Understanding Random Forests: From Theory to Practice. 14077502 Stat

Nakashima K, Ito Y, Yamaguchi-Shinozaki K (2009) Transcriptional Regulatory Networks in Response to Abiotic Stresses in Arabidopsis and Grasses: Figure 1. Plant Physiol 149: 88–95

O’Donnell PJ, Calvert C, Atzorn R, Wasternack C, Leyser HMO, Bowles DJ (1996) Ethylene as a Signal Mediating the Wound Response of Tomato Plants. Science 274: 1914–1917

O’Malley RC, Huang SC, Song L, Lewsey MG, Bartlett A, Nery JR, Galli M, Gallavotti A, Ecker JR (2016) Cistrome and Epicistrome Features Shape the Regulatory DNA Landscape. Cell 165: 1280–1292

Pedregosa F, Varoquaux G, Gramfort A, Michel V, Thirion B, Grisel O, Blondel M, Prettenhofer P, Weiss R, Dubourg V, et al (2011) Scikit-learn: Machine Learning in Python. J Mach Learn Res 12: 2825–2830

Ritchie ME, Phipson B, Wu D, Hu Y, Law CW, Shi W, Smyth GK (2015) limma powers differential expression analyses for RNA-sequencing and microarray studies. Nucleic Acids Res 43: e47–e47

Rojo E, León J, Sánchez-Serrano JJ (1999) Cross-talk between wound signalling pathways determines local versus systemic gene expression in Arabidopsis thaliana: Alternative wound signalling pathways in Arabidopsis. Plant J 20: 135–142

Rose AB, Elfersi T, Parra G, Korf I (2008) Promoter-Proximal Introns in *Arabidopsis thaliana* Are Enriched in Dispersed Signals that Elevate Gene Expression. Plant Cell 20: 543–551

Tierens KFM-J, Thomma BPHJ, Brouwer M, Schmidt J, Kistner K, Porzel A, Mauch-Mani B, Cammue BPA, Broekaert WF (2001) Study of the Role of Antimicrobial Glucosinolate-Derived Isothiocyanates in Resistance of Arabidopsis to Microbial Pathogens. Plant Physiol 125: 1688–1699

Walley JW, Coughlan S, Hudson ME, Covington MF, Kaspi R, Banu G, Harmer SL, Dehesh K (2007) Mechanical Stress Induces Biotic and Abiotic Stress Responses via a Novel cis-Element. PLoS Genet 3: 13

Weirauch MT, Yang A, Albu M, Cote AG, Montenegro-Montero A, Drewe P, Najafabadi HS, Lambert SA, Mann I, Cook K, et al (2014) Determination and Inference of Eukaryotic Transcription Factor Sequence Specificity. Cell 158: 1431–1443

Wilson TJ, Lai L, Ban Y, Steven XG (2012) Identification of metagenes and their interactions through large-scale analysis of Arabidopsis gene expression data. BMC Genomics 13: 237

Yamamoto YY, Yoshioka Y, Hyakumachi M, Maruyama K, Yamaguchi-Shinozaki K, Tokizawa M, Koyama H (2011) Prediction of transcriptional regulatory elements for plant hormone responses based on microarray data. BMC Plant Biol 11: 39

Yan X, Chen S (2007) Regulation of plant glucosinolate metabolism. Planta 226: 1343–1352

Zhang W, Zhang T, Wu Y, Jiang J (2012) Genome-Wide Identification of Regulatory DNA Elements and Protein-Binding Footprints Using Signatures of Open Chromatin in *Arabidopsis*. Plant Cell 24: 2719–2731

Zou C, Sun K, Mackaluso JD, Seddon AE, Jin R, Thomashow MF, Shiu S-H (2011) Cisregulatory code of stress-responsive transcription in Arabidopsis thaliana. Proc Natl Acad Sci 108: 14992–14997

