## Supplemental figures for "Modeling gene regulation in response to wounding: temporal variations, hormonal variations, and specialized metabolism pathways induced by wounding"

### Supplemental Figure 1

**A.**

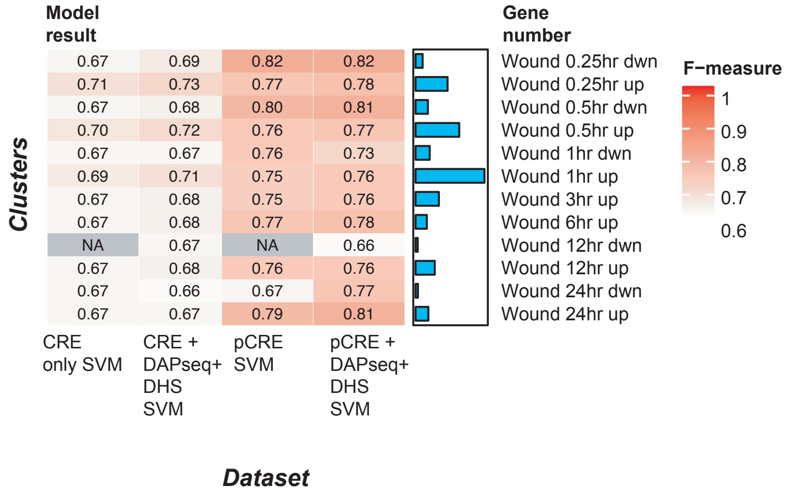


**B.**

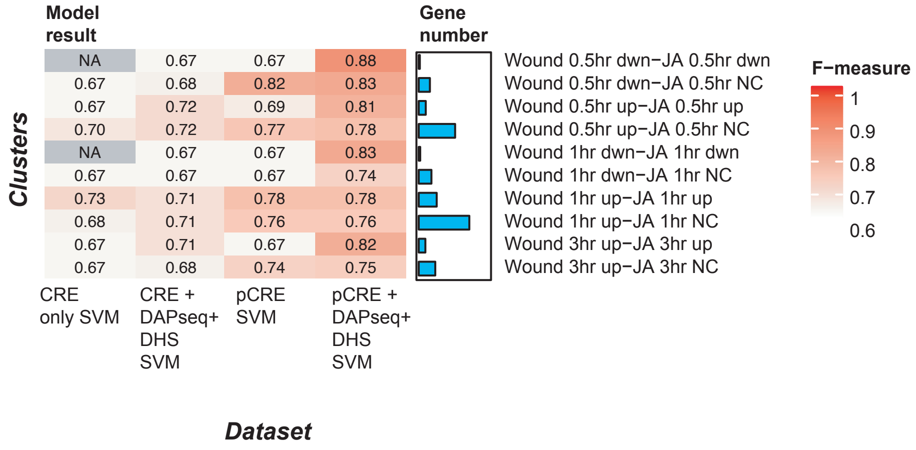

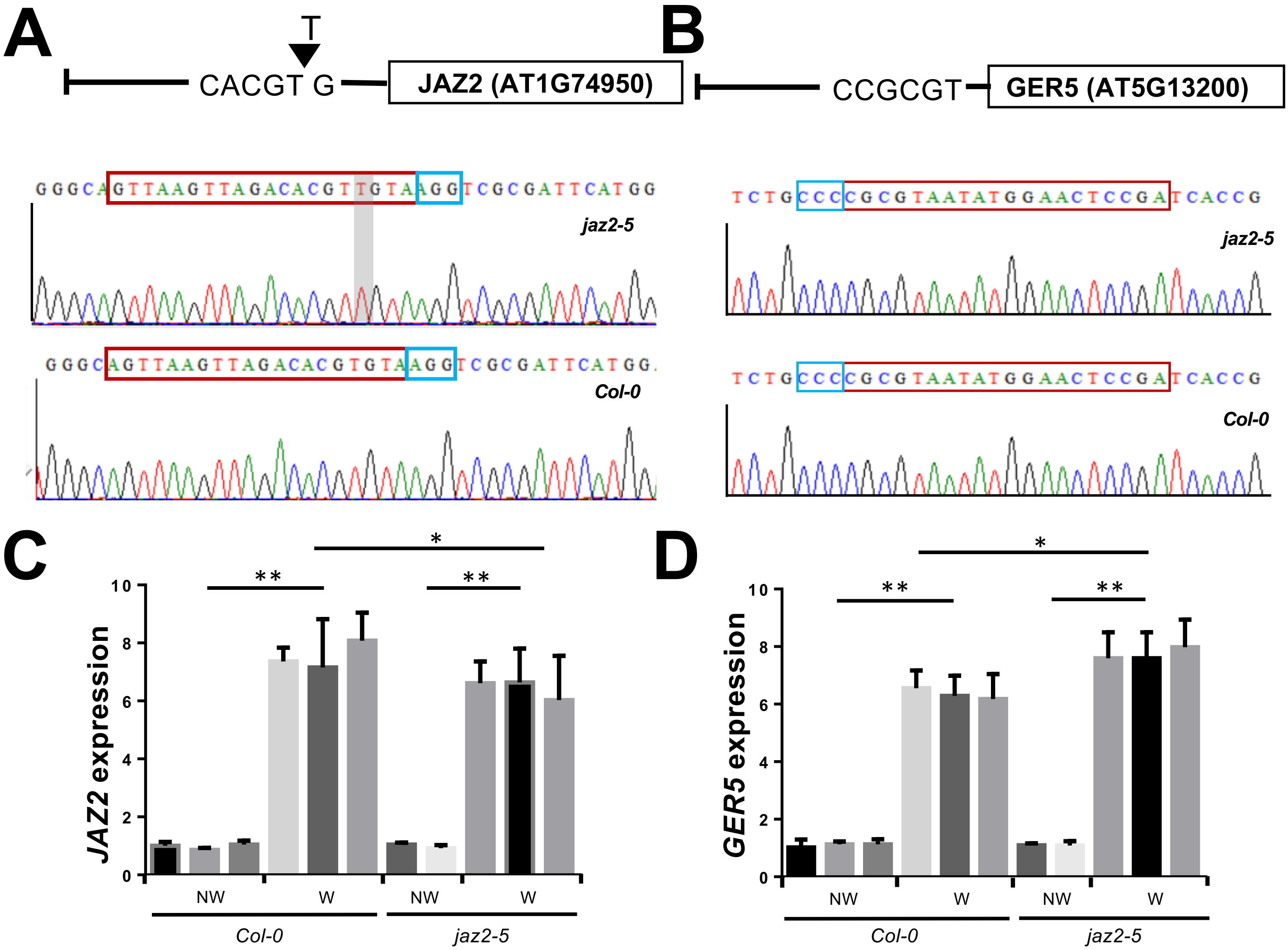


**Supplemental Figure 2. Mutation in CACGTG motif of *JAZ2* promoter led to down-regulation of *JAZ2* expression following wound treatment.** A. CRISPR/Cas9-mediated mutation in CACGTG motif of *JAZ2* promoter region in *jaz2-5* mutant. B. No mutation in CCGCGT motif of *GER5* promoter in the *jaz2-5*. Note that the CRISPR/Cas9 against the *GER5* promoter did not result in nucleotide changes in the region. Upper chromatogram shows the CRISPR/Cas9-targeted promoter regions of the *JAZ2* and *GER5* promoter. Lower chromatogram displays its corresponding region in Col-0. The T insertion within CACGTG motif of the *JAZ2* promoter was marked with shadow on the chromatogram. Blue and red boxes highlight PAM sequence and gRNA target region, respectively. C. Wound-induced *JAZ2* expression in Col-0 and *jaz2-5*. D. Wound-induced *GER5* expression in Col-0 and *jaz2-5*. The transcript abundances of *JAZ2* or *GER5* were normalized to *ACTIN2*. NW indicates no wound treatment and W indicates wound treatment. Three biological replicates are individually demonstrated with error bars, obtained from three technical repeats. One-way analysis of variance (ANOVA) was applied to analyze statistical significances of the gene expression levels, and the differences are indicated by asterisks. (* $p < 0.05$, ** $p < 0.01$)

#### Supplemental Figure 3

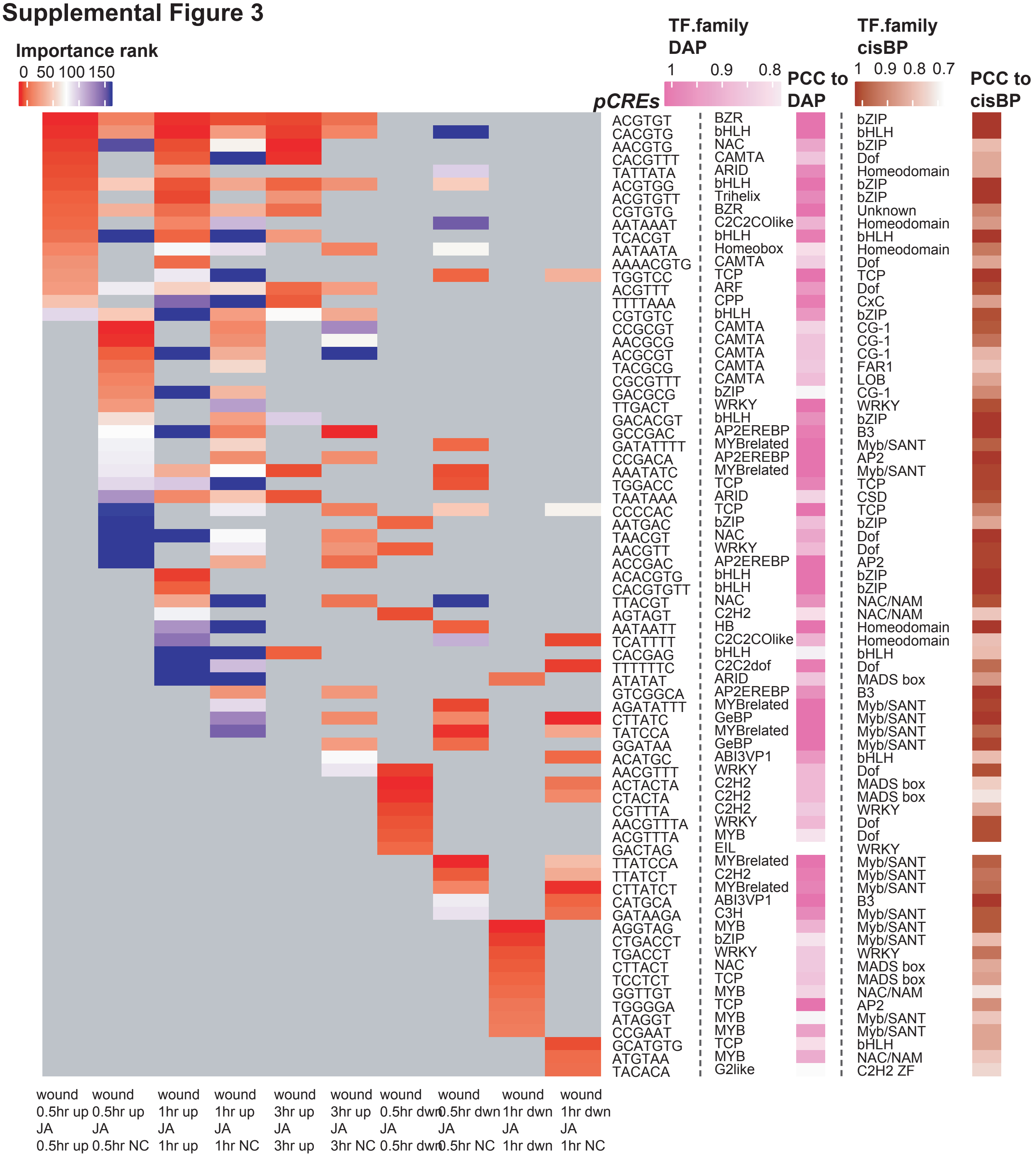
